# Integrated single cell analysis shows chronic alcohol drinking disrupts monocyte differentiation in the bone marrow niche

**DOI:** 10.1101/2023.03.29.534727

**Authors:** Sloan A. Lewis, Brianna M Doratt, Qi Qiao, Madison B. Blanton, Kathleen A. Grant, Ilhem Messaoudi

## Abstract

Chronic alcohol drinking rewires circulating monocytes and tissue-resident macrophages towards heightened inflammatory states with compromised anti-microbial defenses. As these effects remain consistent in short-lived monocytes after a 1-month abstinence period it is unclear whether these changes are restricted to the periphery or mediated through alterations in the progenitor niche. To test this hypothesis, we profiled monocytes/macrophages and hematopoietic stem cell progenitors (HSCP) of the bone marrow compartment from rhesus macaques after 12 months of ethanol consumption using a combination of functional assays and single cell genomics. Bone marrow-resident monocytes/macrophages from ethanol-consuming animals exhibited heightened inflammation. Differentiation of HSCP *in vitro* revealed skewing towards monocytes expressing neutrophil-like markers with heightened inflammatory responses to bacterial agonists. Single cell transcriptional analysis of HSCPs showed reduced proliferation but increased inflammatory markers in mature myeloid progenitors. We observed transcriptional signatures associated with increased oxidative and cellular stress as well as oxidative phosphorylation in immature and mature myeloid progenitors. Single cell analysis of the chromatin landscape showed altered drivers of differentiation in monocytes and progenitors. Collectively, these data indicate that chronic ethanol drinking results in remodeling of the transcriptional and epigenetic landscapes of the bone marrow compartment leading to altered functions in the periphery.

## INTRODUCTION

Alcohol drinking is widespread with more than 2 billion current drinkers worldwide (1). Alcohol and its metabolic products induce organ damage and increase incidence of cardiovascular disease (2, 3), several types of cancer (4–7), liver cirrhosis (8), and sepsis (9). Moreover, heavy alcohol drinking has been linked to increased susceptibility to several bacterial and viral infections (10–13). Increased vulnerability to infectious diseases is believed to be mediated in part by functional, transcriptomic, and epigenomic changes in blood monocytes and tissue-resident macrophages leading to increased inflammation but compromised antimicrobial responses (14–17). Whether these changes are limited to the peripheral myeloid compartments or can be traced to progenitor cells in the bone marrow has yet to be determined.

Monocytes continuously arise from hematopoietic stem cell progenitors (HSCP) in the bone marrow through progressively restricted lineage committed progenitors (18, 19). Two independent pathways of monocyte production have been reported in mice starting from common myeloid progenitors (CMP) and proceeding through either granulocyte-monocyte progenitors (GMP) to monocyte progenitors (MP) or monocyte-DC progenitors (MDP) to common monocyte progenitors (cMoP) (19, 20). Myelopoiesis in humans is not as well characterized, but CMP, GMP, MDP, and cMoP populations have been identified (18, 21, 22). Mature monocytes can be further classified into three subsets, classical, intermediate, and non-classical, which can be found in the bone marrow as well as in circulation (23). Infection, inflammation, or other stress factors can alter monocyte production and even induce “emergency monopoiesis” from the bone marrow compartment (19, 24, 25).

Chronic heavy drinking (CHD) is known to affect bone marrow stem cells and hematopoiesis. Specifically, lymphopenia, anemia, and thrombocytopenia are observed in patients with alcohol use disorder (26–31). Studies in rodent models of alcohol exposure have reported impairment of hematopoietic precursor cell activation as well as perturbation of granulocyte precursor responses and differentiation resulting in reduced bacterial clearance (32, 33). Studies in non-human primates (NHP) have reported impaired mitochondrial function of HSCP and alterations in the bone marrow niche with chronic ethanol consumption (34). Finally, alcohol use and simian immune deficiency virus (SIV) co-infection results in increased numbers of mature macrophage and osteoclasts in the bone marrow with (35). The impact of alcohol drinking on monopoiesis in the bone marrow has not been examined.

In this study, we utilize an NHP model of voluntary ethanol consumption to assess the impacts of CHD on monocytes and their progenitors in the bone marrow compartment. We collected bone marrow cells from the femurs of male and female animals engaged in CHD (defined as average daily consumption of >3g ethanol per kilogram body weight) for 12 months. We performed phenotypic, functional, and single cell transcriptomic/epigenomic assays on both CD14+ monocytes and CD34+ progenitor cells within the bone marrow. Our analysis revealed broad increases in inflammatory and oxidative stress-associated signatures in bone marrow CD34+ and CD14+ cells with CHD. This was accompanied by skewing of the CD34+ cells to produce more neutrophil-like monocytes with heightened inflammatory properties as well as increased inflammation in intermediate monocytes in the bone marrow. Finally, these functional and transcriptional changes were accompanied by epigenetic changes in monocyte progenitors.

## RESULTS

### CHD-induced heightened inflammatory state of circulating monocytes is sustained through 1-month abstinence period

We have previously shown that CHD in NHPs leads to heightened inflammation in circulating monocytes and tissue resident macrophages that is mediated by epigenetic and transcriptional changes (14, 36). As monocytes are relatively short-lived cells (5-7 days in circulation), we set to determine whether circulating monocytes would revert to a control state following a 1-month abstinence period (23). We collected peripheral blood mononuclear cells (PBMC) from male macaques under a repeated abstinence protocol and terminated following the final 1-month abstinence period (n=4) and control animals (n=3) (37) (**Figure 1A, B and Supp. Table 1**). We have previously reported increased percentages of monocytes in circulation as well as heightened TNF production in response to lipopolysaccharide (LPS) stimulation from these animals (14). Following a 1-month abstinence period, the frequency of circulating monocytes remained increased (**Figure 1C**) as was the frequency of TNF – secreting monocytes and the concentration of TNF in response to LPS (**Figure 1C**).

**Figure 1:**
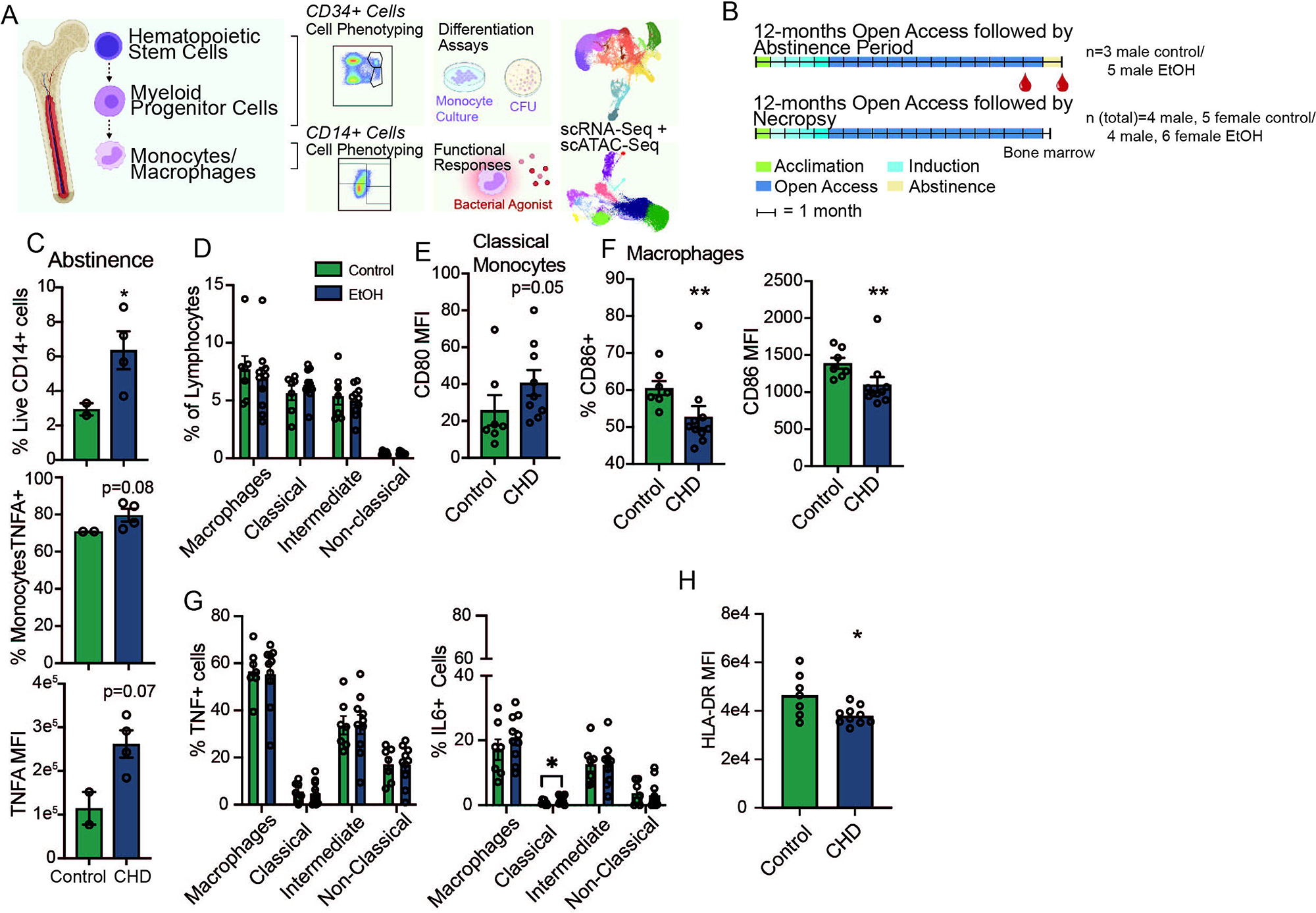
Inflammatory blood monocyte phenotype with CHD extends through abstinence. A) Experimental design for study partially created on Biorender.com. B) CHD timeline (+/-abstinence) and blood/bone marrow collection for macaque cohorts. C) Flow cytometry phenotyping and intracellular cytokine staining (ICS) after 16-hour LPS stimulation were performed on total PBMC after 1-month abstinence. Bar plots showing % live CD14+ (top), % TNFα + monocytes (middle), and TNFα MFI from monocytes. ICS measurements were corrected for the unstimulated condition. D) Percentages of classical, intermediate, and non-classical monocytes, and CD169+ macrophages from live cells. E) Bar plot of CD80 MFI on classical monocytes. F) Bar plots of the percentage of CD86+ (left) and CD86 MFI (right) on macrophages. G) Total bone marrow cells were stimulated with a bacterial TLR cocktail (Pam3CSK4, LPS, and FSL-1) and the percentage of TNFα+ (left) and IL-6+ (right) were measured in each monocyte and macrophage population and corrected for the unstimulated condition. H) HLA-DR MFI was measured in macrophages after bacterial agonist stimulation by flow cytometry and corrected for the unstimulated condition. Statistical significance was tested by t-test with Welch’s correction where *=p<0.05, **=p<0.01.

### Chronic CHD induced subtle changes in the functional responses of monocytes and macrophages in the bone marrow

The persistence of the inflammatory phenotype of monocytes suggests that CHD may impact the progenitor population in the bone marrow compartment. To test this hypothesis, we profiled monocytes that reside within the bone marrow compartment from female (n=6) and male (n=4) monkeys after 12 months of CHD as well as control animals (n=5 females; n=4 males) (**Figure 1D** **and Supp. Figure 1A**). No differences were detected in the relative abundance of classical, intermediate, and non-classical monocyte and macrophage populations (**Figure 1D**). However, we noted an increase in the level of the co-stimulatory molecule CD80 on classical monocytes as indicated by higher mean fluorescence intensity (MFI) suggesting increased baseline activation (**Figure 1E**). Interestingly, expression of activation marker CD86 on bone marrow macrophages was lower in CHD animals (**Figure 1F**).

To determine if bone marrow monocytes exhibited the same heightened inflammatory responses as circulating monocytes, total bone marrow cells were stimulated with a bacterial TLR cocktail (Pam3CSK4, LPS, and FSL-1) and percentages of TNF and IL-6 producing monocytes/macrophages were determined using flow cytometry (**Figure 1G**). No differences were noted in responses of the intermediate or non-classical monocyte populations; however, a greater percentage of classical monocytes from CHD animals produced IL-6 (**Figure 1G**). Moreover, surface expression of HLA-DR on macrophages from CHD animals was downregulated to a greater extent following stimulation (**Figure 1H**).

### scRNA-Seq of bone marrow monocytes and macrophages reveals increased oxidative stress and inflammatory transcriptional signatures with CHD

To capture transcriptional changes occurring in bone marrow monocytes with CHD at a higher resolution, we performed scRNA-Seq on purified bone marrow CD14+ cells obtained following 12 months of chronic ethanol drinking (n=3 female controls pooled due to low cell numbers, 3 female CHD pooled, 4 male controls, 4 male CHD; **Supp. Figure 1B**). After integrating all datasets, 11 clusters were identified that could be grouped based on the expression of canonical markers into: 8 classical monocyte clusters (C 1-8; *CD14, LYZ*), 1 intermediate monocyte cluster (Int.; *MAMU-DRA, S100A10*), 1 non-classical monocyte cluster (N.C.; *FCGR3*), and 1 macrophage cluster (Mac.; *FABP4, SIGLEC1*) (**Figure 2A,B** **and Supp. Figure 1C**). The 8 clusters within classical monocytes each expressed a unique gene expression profile as identified by the *FindAllMarkers* function in *Seurat* (**Figure 2C** **and Supp. Table 2**). Chronic CHD was associated with a significant increase in classical cluster C8 (**Figure 2D**), which was defined by high expression of *CREBRF, HERPUD1*, *FOSB* (**Figure 2C**). *CXCR4*, the receptor for SCF-1 and shown to be expressed on a population of pro-monocytes (38) was also highly expressed in this cluster (**Figure 2E**).

**Figure 2:**
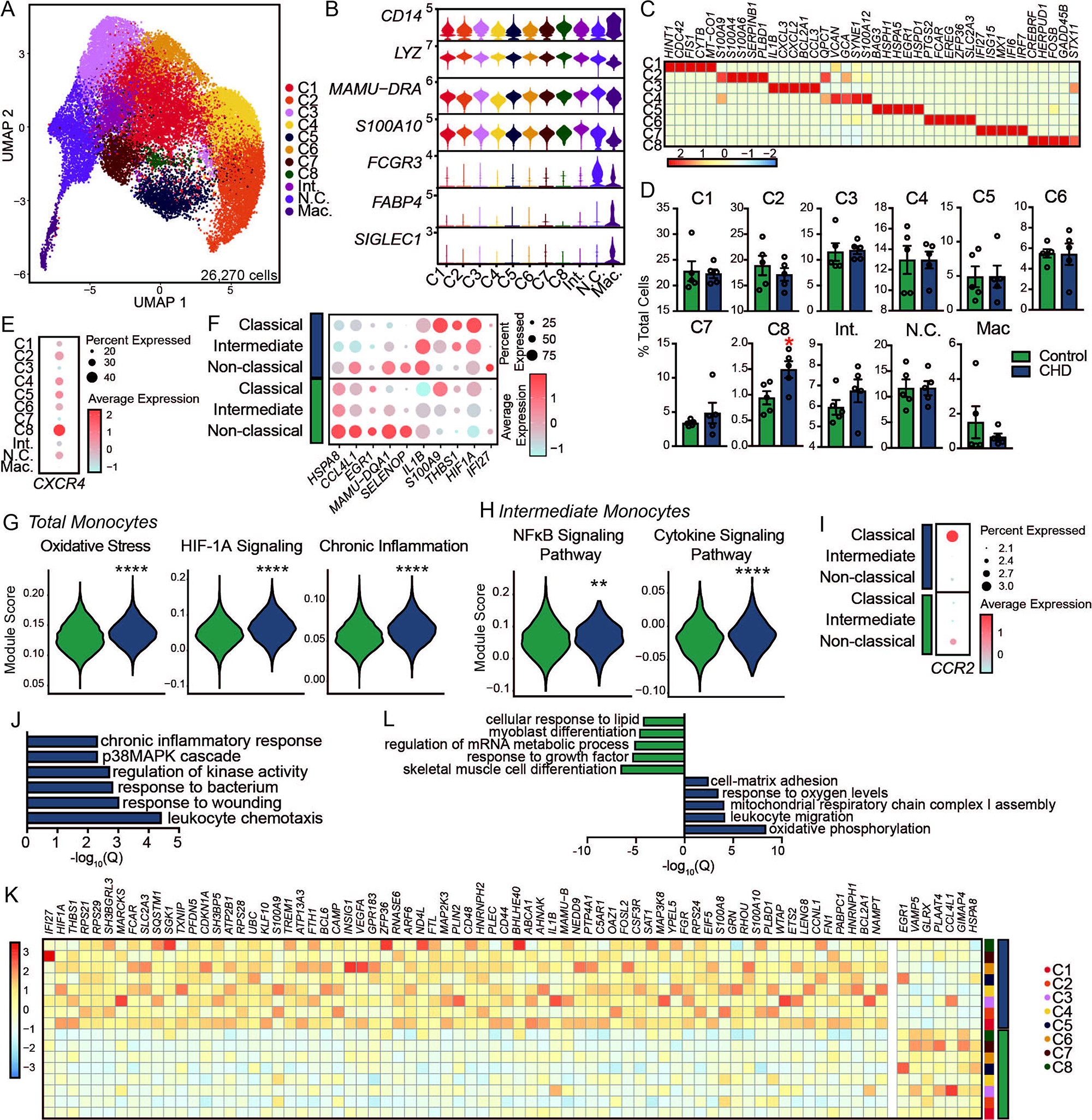
Shift in the single cell transcriptional profiles of CD14+ cells from the bone marrow of macaques with CHD A) UMAP clustering of 26,270 cells. B) Stacked violin plot showing expression of genes identified using Seurat’s *FindAllMarkers* function. C) Heatmap showing averaged marker gene expression of highly expressed genes from each classical monocyte cluster. D) Bar plots showing percentage of total cells contributing to each monocyte cluster. E) Dot plot showing expression of *CXCR4* across each cluster where the size of the dot represents percent of cells expressing the gene and the color represents an averaged expression value. F) Dot plot showing expression of up-and down-regulated DEG with CHD common to all three monocyte subsets where the size of the dot represents percent of cells expressing the gene and the color represents an averaged expression value. G) Violin plots representing module score expression for oxidative stress, HIF-1α signaling, and chronic inflammation pathways in total monocytes. H) Violin plots representing module score expression for NFκB and cytokine signaling pathways in intermediate monocytes. Statistical significance was tested using Mann-Whitney test. I) Dot plot showing expression of *CCR2* across each monocyte subset split by CHD and control where the size of the dot represents percent of cells expressing the gene and the color represents an averaged expression value. J) Bar plot representing -log_10_(q-value) functional enrichment scores for genes upregulated in classical monocyte clusters with CHD. K) Heatmap showing averaged gene expression of DEG from each classical monocyte cluster split by CHD and control groups. L) Bar plot representing -log_10_(q-value) functional enrichment scores for genes up- and downregulated in the macrophage cluster with CHD.

It has been suggested that there are two distinct lineages of monocyte production in the bone marrow (18–20); therefore, we assessed the impact of CHD on the frequency of MDP-derived versus GMP-derived classical monocytes. C2 and C4 were classified as GMP-derived based on high levels of *S100A8/9* while C1, C3, C5, C6, C7, C8 were classified as MDP-derived based on expression of *CD74* and *MAMU-DRA* (**Supp. Figure 1D**). While we did not note significant changes in the distributions of MDP-(C1,3-8) versus GMP-derived (C2,4) clusters with CHD (**Figure 2D**), the GMP-derived clusters had higher expression of *S100A8/9* and the MDP-derived clusters had lower expression of *CD74/MAMU-DRA* with CHD (**Supp. Figure 1D**).

We next set out to uncover the impact of CHD on the transcriptional profile of bone marrow resident monocytes. Broadly, we noted decreased expression of *HSPA8, CCL4L1, EGR1, MAMU-DQA1*, and *SELENOP* and increased expression of *IL1B, S100A9, THBS1, HIF1A*, and *IFI27* across classical, intermediate, and non-classical clusters with CHD (**Figure 2F**). This was accompanied by increased module scores associated with oxidative stress, HIF1A Signaling, and chronic inflammation in all monocyte clusters with CHD (**Figure 2G**). Increased scores of modules associated with NFκB and cytokine signaling pathways were also increased within the intermediate cluster with CHD (**Figure 2H** **and Supp. Table 3**). Interestingly, expression of *CCR2*, a marker associated with monocyte egress from the bone marrow compartment (39), was highest on non-classical monocytes from controls but on classical monocytes from the CHD group, indicating potential dysregulation of monocyte export from bone marrow with alcohol (**Figure 2I**). Differential gene expression analysis within the classical clusters revealed significant upregulation of genes mapping to chemotaxis (*CSF3R, VEGFA*), response to wounding (*CD44, FN1*), and chronic inflammatory response (*S100A8, S100A9*) with CHD (**Figure 2J,K**). Within the non-classical subset, expression of genes important for defense response (*MRC1, STAT1*) and stem cell differentiation (*MEF2C, CITED2*) was reduced, while that of genes that play a role in wound healing (*SERPINA1, THBS1*) and migration (*MIF, FN1*) was increased with CHD (**Supp. Figure 1E**).

Bone marrow macrophages have critical functions in homeostatic maintenance of stem cells and bone marrow niche (40, 41). Although the frequency of this cluster did not differ between the two groups (**Figure 2D**), considerable transcriptional changes were detected with CHD. DEG downregulated with CHD mapped to GO terms associated with cellular responses to stimuli (*CCL8, CCL13, CCL24, CCL4L1*) and developmental processes (*DDX5, EGR1, FOS, BTG2*) (**Figure 2L** **and Supp. Figure 1F**). On the other hand, DEG upregulated with CHD mapped to mitochondrial respiration (*MT-ATP8, MT-CO2, MT-ND2, NDUFA1*) and leukocyte migration processes (*IFI27, TREM1, VEGFA, VCAN*) (**Figure 2L** **and Supp. Figure 1G**). These data suggest CHD alters the transcriptional landscape of bone marrow-resident monocytes and macrophages, with the potential to impact bone marrow niche homeostasis, monocyte export, and responses to external stimuli.

### CHD skews monocyte differentiation from CD34+ progenitors

As monocytes are being constantly produced by and stored in the bone marrow, we investigated the impact of CHD on the ability of CD34+ progenitor cells to differentiate into monocytes. To that end, CD34+ cells were purified and cultured for 7 days in the presence of a monocyte skewing supplement (**Figure 3A**). Cultures from CHD animals showed reduced capacity to differentiate into CD14+CD34-cells compared to control animals with more cells retaining CD34+ expression (**Figure 3B-C**). Expression of monocyte maturation markers CD115 and CD11C increased similarly in the cultured CD14+ populations regardless of CHD exposure (**Supp. Figure 2A**).

**Figure 3:**
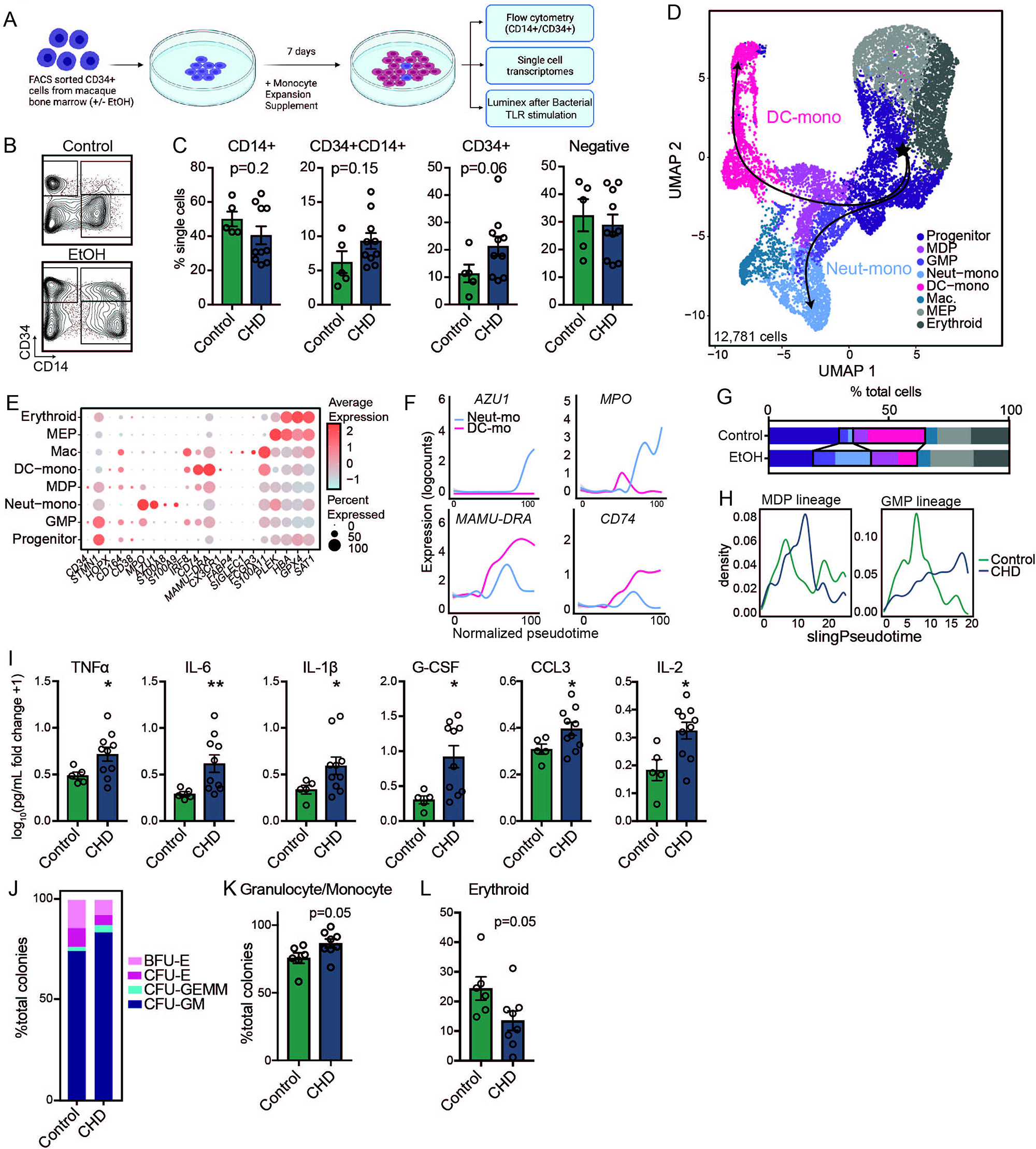
CHD alters CD34+ progenitor cell differentiation to monocytes. A) Experimental design for this figure created on Biorender.com. Sorted CD34+ cells from control and CHD macaque bone marrow were cultured in monocyte differentiation media supplement for 7 days. B) Example flow gating showing CD34+ versus CD14+ cells. C) Bar plots showing quantification of the culture output by flow cytometry. D) The same cultures were pooled from each group and subjected to 10X scRNA-Seq. UMAP projection of 12,781 cells overlayed with Slingshot pseudotime lineage lines. E) Dot plot showing expression of genes identified using Seurat’s *FindAllMarkers* function across each cluster where the size of the dot represents percent of cells expressing the gene and the color represents an averaged expression value. F) Log expression of *AZU1, MPO, MAMU-DRA, CD74* plotted for each cell across the indicated scaled Slingshot pseudotime trajectory (trendline shown). G) Bar plots showing representative percentages of each cluster across control and CHD groups. H) Cell density plots for Control and CHD groups across each trajectory lineage determined by Slingshot. I) Cultures were stimulated for 6 hours with a bacterial TLR cocktail (Pam3CSK4, LPS, and FSL-1). Bar plots showing the log_10_(fold change +1) concentration of each of the indicated analytes measured by Luminex. J) Stacked bar plot showing percentages of CFU-GM, CFU-GEMM, CFU-E, and BFU-E colonies. (K-L) Bar plots showing the percentage of Granulocyte/Monocyte and Erythroid colonies from total colonies across control and CHD groups.

To further assess differences in monocyte differentiation between the control and CHD groups, we performed scRNA-Seq on the differentiated cells (**Figure 3D**). Using highly expressed gene markers, we were able to identify progenitor cells (*CD34, STMN1, CD38*), GMP-derived monocytes (*S100A8/9*), MDP-derived monocytes (*IRF8, CX3CR1*), monocyte-derived macrophages (*SIGLEC1, S100A11*), and two clusters of megakaryocyte/erythroid progenitors (*PLEK, HBA, GPX4*) (**Figures 3D,E** **and Supp. Table 2**). Pseudotime analysis identified a GMP lineage defined by increasing expression of *AZU1* and *MPO*, and an MDP lineage defined by increasing expression of *MAMU-DRA* and *CD74* (**Figure 3F**). While CD34+ cells from control animals differentiated primarily along the MDP lineage into DC-monocytes, CD34+ cells from the CHD group differentiated along the GMP lineage towards Neutrophil-monocytes (**Figure 3G,H**). Next, differentiated CD34+ cells were stimulated with a bacterial TLR cocktail and cytokine, chemokine, and growth factor production was measured by Luminex (**Figure 3A**). Production of TNFα, IL-6, IL1β, G-CSF, CCL3 (MIP-1α) and IL-2 was significantly increased in CHD cultures after the stimulation suggesting that CD34+ progenitor cells from CHD animals are poised towards a heightened inflammatory response (**Figure 3I**).

Finally, to assess the impact of CHD consumption on the ability of CD34+ progenitor cells to differentiate into cells of the myeloid lineage, we performed a colony forming unit (CFU) assay. No differences were noted in total number of colonies after 7 or 10 days of culture **(Supp. Figure 2B)**, but a skewing of the progenitors towards granulocyte/monocyte-containing colonies (CFU-GM and CFU-GEMM) and away from erythroid only colonies (CFU-E and BFU-E) was evident at day 10 in CHD cultures (**Figure 3J-L**). These observations indicate that CHD alters the differentiation potential of CD34+ progenitors, skewing monocyte differentiation towards a more inflammatory GMP-derived lineage and increasing the production of granulocyte/monocyte progenitor colonies.

### scRNA-Seq of CD34+ progenitors reveals reduced proliferation but increased oxidative stress and inflammatory pathways in myeloid progenitors with CHD

Next, we assessed the impact of chronic ethanol consumption on the distribution of major progenitor populations by flow cytometry (**Supp. Figure 2C**). Frequencies of CD34+CD38-CD45RA-CD90+ HSCP, and CD34+CD38+CD45RA-CD123+CD64-CMP were modestly increased in the CHD group whilst those of CD34+CD38-CD45RA-CD90-multipotent progenitors (MPP) were significantly increased (**Supp. Figure 2D**). To further interrogate the impact of chronic ethanol consumption on these progenitors, we profiled purified CD34+ bone marrow cells by scRNA-Seq (n=3 female controls, 4 male controls, 3 female CHD, 4 male CHD). UMAP clustering revealed 17 clusters based on highly expressed gene markers (**Supp.** **Figure 3A,B** **and Supp. Table 2)**. HSC clusters expressed high levels of *HOPX, CD164, MKI67*, and *STMN1* while cells in the erythroid lineage including megakaryocyte/erythroid progenitors expressed *CPA3* and *KLF1* and those from the lymphoid lineage expressed *RAG1, IL7R*, and *CD79B* (**Supp. Figure 3C)**. Within the myeloid lineage, MDP could be identified based on the expression of *IRF8, MAMU-DRA,* and *BATF3,* while GMP were identified based on the expression of *LYZ, ELANE*, and *MPO* (**Supp. Figure 3C**). Identification of CMP, GMP and MDP subsets was further confirmed using module scoring based on gene lists from the Human Cell Atlas bone marrow single cell dataset (42) (**Supp. Figure 3D and Supp. Table 3**). Pro-neutrophils expressed lower levels of *LYZ* and high levels of *ELANE* and *MPO* (**Supp. Figure 3C**). More differentiated monocyte clusters (cMoP, MP, and pro-monocytes) were defined by increased expression of *FCER1A* and *CD14* (**Supp. Figure 3C**). CHD was associated with modest reductions in the CMP/GMP and GMP-2 clusters that was accompanied by a slight increase in the CLP/pre T cell clusters (**Supp. Figure 3E**).

The myeloid subsets were further subclustered for a more targeted analysis of the effects of CHD on this lineage. Further, contaminating mature bone marrow monocytes were removed using the CD14+ scRNA-Seq data described in Figure 2 (**Supp. Figure 4A**). The progenitor CD34+ cells were clustered and annotated according to the expression of myeloid lineage marker genes into pro-neutrophils (*ELANE, MPO, AZU1*), monocyte progenitors (*LYZ, S100A8/9*), and pro-DC (*FCER1A, IRF8*) (**Figure 4A,B**). CHD was associated with significantly reduced frequency of the proliferating MP/GMP cluster (**Figure 4C**). Slingshot trajectory analysis identified three major lineage paths (**Figure 4A**) that were delineated by expression of genes associated with pro-DC, pro neutrophil, and pro-monocyte development and were relatively similar in density between control and CHD groups (**Figure 4D,E** **and Supp. Figure 4B,C**).

**Figure 4:**
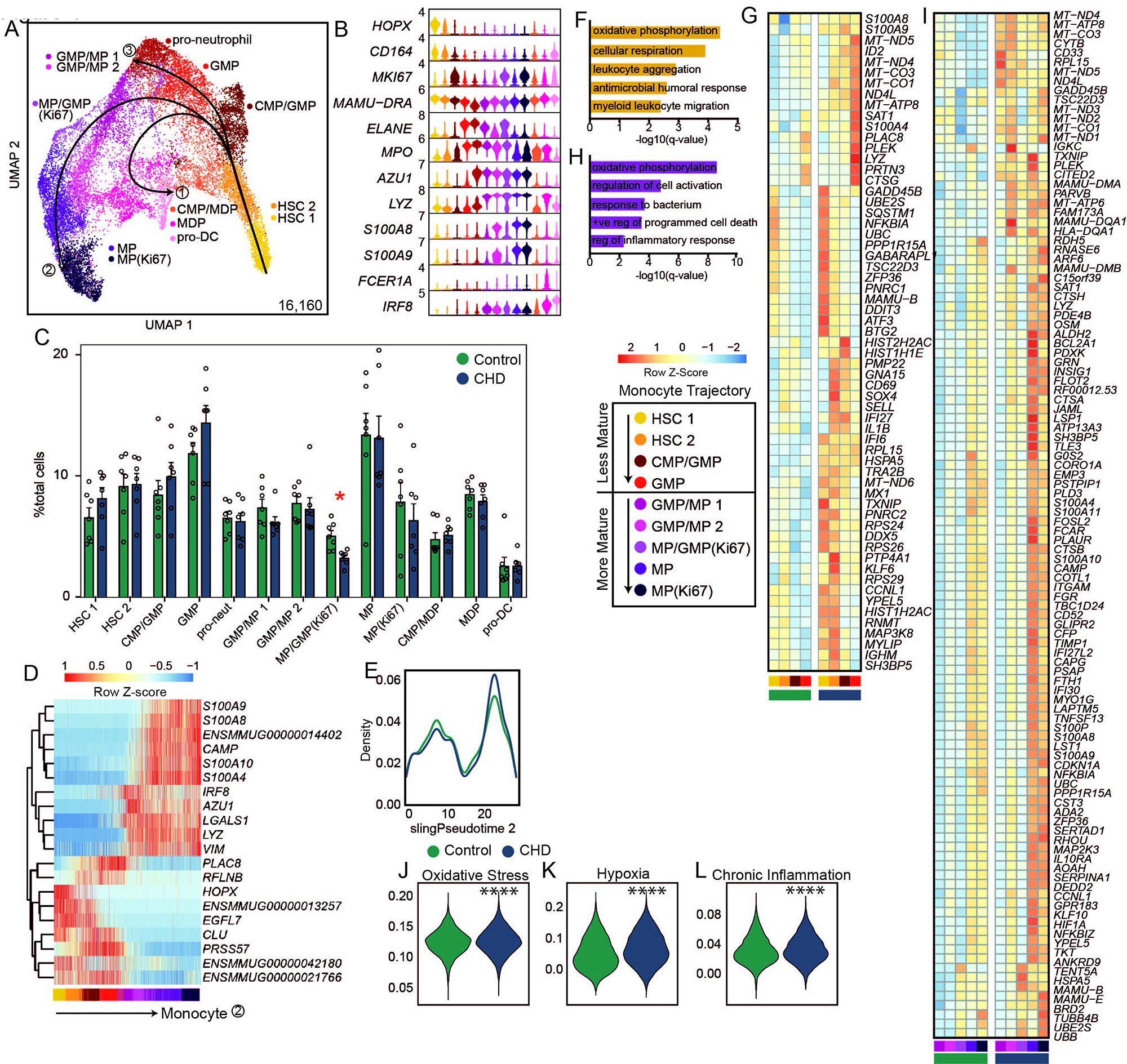
CHD CD34+ bone marrow myeloid progenitor single cell transcriptional profiles A) Myeloid lineage UMAP (16,160 cells) with Slingshot lineage projection lines. B) Stacked violin plot of marker gene expression. C) Cluster percentages between CHD and control groups. Compared by two-way ANOVA with multiple comparisons. D) Heatmap of genes explaining the monocyte lineage trajectory. E) Cell density plot for Control and CHD groups across the monocyte trajectory lineage determined by Slingshot. F,H) Functional enrichment terms from less (F) and more (H) mature clusters. G,I) Averaged gene expression of upregulated DEG from less (G) and more (I) mature clusters split by CHD and control groups. J-L) Violin plots for indicated module scores across groups. Statistical analysis was performed by Mann-Whitney. Unless indicated, statistical significance was tested by t-test with Welch’s correction. *=p<0.05, ****=p<0.0001.

We next carried out differential gene expression grouped by less mature (HSC 1, HSC 2, CMP/GMP, GMP) and more mature (GMP/MP 1, GMP/MP 2, MP/GMP (Ki67), MP, MP (Ki67)) monocyte progenitor cells identified based on the pseudotime analysis (**Figure 4D**). In the less mature progenitor cells, CHD consumption led to increased expression of genes involved in “oxidative phosphorylation” and “leukocyte aggregation” processes (**Figure 4F,G**), but a downregulation of genes involved in “cellular oxidant detoxification” and “regulation of chemokine production” processes (**Supp. Figure 4D,E**). Similarly, expression of genes enriching to “oxidative phosphorylation” and “regulation of cell activation” processes was increased in the more mature progenitors with CHD consumption (**Figure 4H,I**) while genes enriching to “cytoplasmic translation” and “peptide biosynthetic process” were downregulated with CHD (**Supp. Figure 4F,G**). Finally, module scores of oxidative stress, hypoxia, and chronic inflammation were significantly upregulated in all progenitor clusters with CHD consumption (**Figure 4J-L**). These data suggest that there are transcriptional changes in the CD34+ compartment with CHD that point to increased inflammation and oxidative stress. This is further evidence that the impact of CHD on monocytes is occurring at the progenitor level.

### scATAC-Seq of CD34+ and CD14+ cells reveal changes in the epigenome of monocyte progenitors

Finally, to determine whether epigenetic changes were driving transcriptional changes within progenitor cells and bone marrow resident monocytes, we sorted and pooled CD14+ and CD34+ cells from the bone marrow and performed single cell ATAC sequencing (n=3 male controls, 2 female CHD, 2 male CHD). Initial UMAP clustering revealed 13 clusters, which we defined based on gene scores of canonical lineage markers (**Supp. Figures 5A-C and Supp. Table 4**). We then extracted HSC, CMP-GMP, and CD14+ subsets based on expression of the canonical markers for these subsets (*HPOX8, IRF78, ELANE, MPO, LYZ, MAMU-DRA, CD14*) and re-clustered them (**Figure 5A**). This analysis revealed 6 clusters (**Figure 5A**) defined by genes scores: HSC (*HOPX*), GMP 1 and 2 (*ELANE*), MP/GMP (*LYZ*), cMoP (LYZ), and monocytes (*CD14*) (**Figure 5B**). Cluster annotation was further confirmed by transcription factor (TF) motif deviation scores that showed monocytes and more mature progenitors contained a greater abundance of motif binding sites for FOSL2 and CEBPA, whereas HSC contained a greater abundance of motif binding sites for HOXB8 and JUNB (**Supp. Figure 5D**).

**Figure 5:**
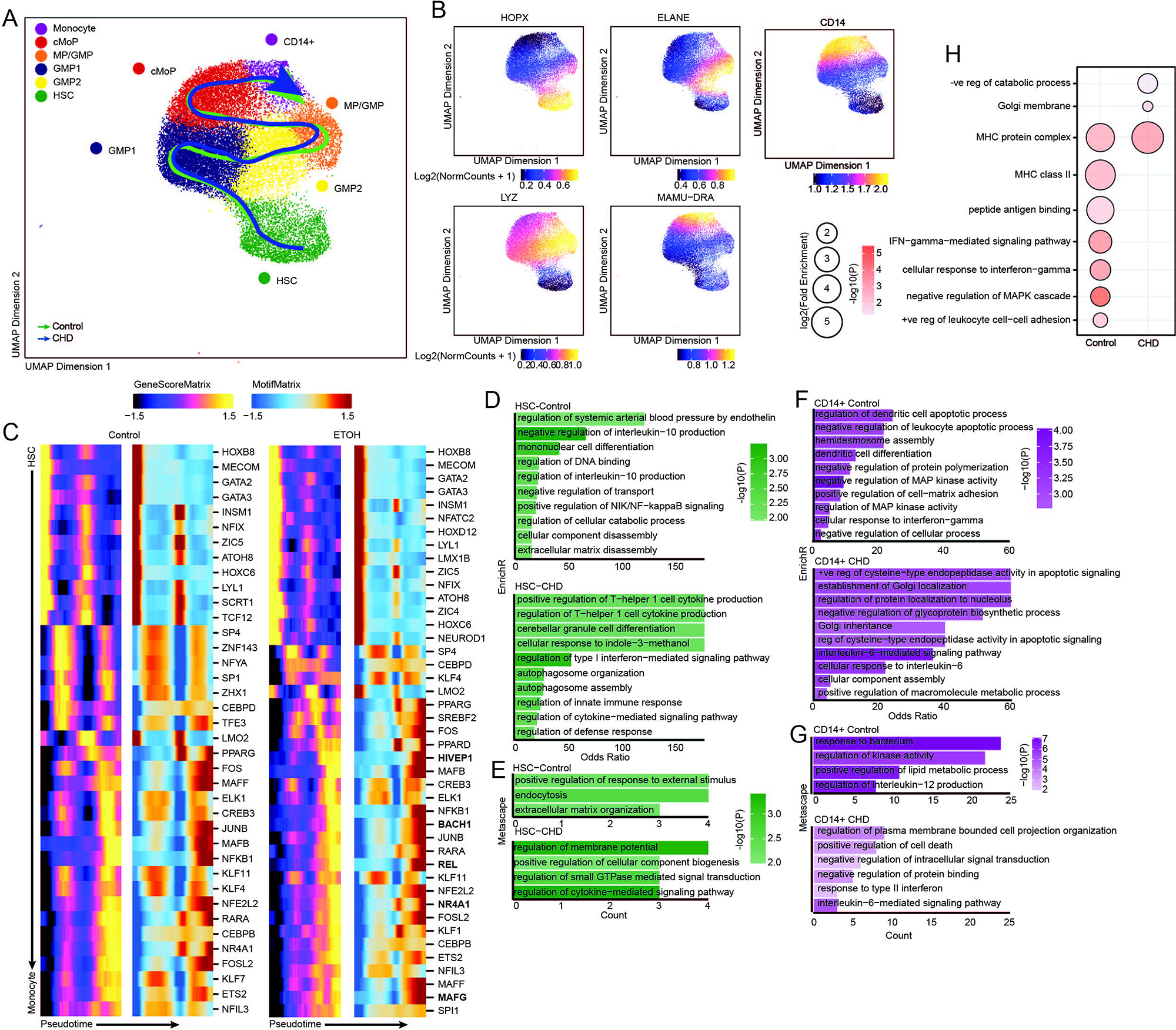
CD14+ and CD34+ bone marrow cell scATAC-Seq. A) UMAP projection of myeloid lineage cells based on gene accessibility with pseudotime lineage lines for control and CHD groups. B) Feature plots of marker genes from the gene score matrix. C) Integrative analysis of TF gene scores and motif accessibility across the monocyte lineage pseudotime in control and CHD cells. Genes listed on CHD heatmap are observed only with CHD. D-G) Functional enrichment from EnrichR (D,F) and Metascape (E,G) databases for differentially accessible regions more open in the indicated group for HSC (D-E) and pro-monocyte (F-G) clusters. Color indicates the -log10(P) value and length indicates the Odds Ratio from EnrichR (D,F) or the number of genes mapping to the term from metascape (E,G). (H) Functional enrichment determined by GO Biological Process within GREAT database for differentially accessible regions more open control (left) and CHD (right) groups for the pro-monocyte cluster. Color indicates the -log10(P) value and length indicates the log2(Fold Enrichment).

As the frequency of cell subsets were not changed with CHD **(Supp. Figure 5E**), we performed trajectory analysis using Slingshot to uncover potential differences in monocyte differentiation between the two groups (**Figure 5A,C**). CHD induced epigenetic changes associated with drivers of differentiation to monocytes where motif accessibility for key TFs including MAFG, HIVEP1, REL, NR4A3, and BACH1 was increased with CHD (**Figure 5C**). Next, we assessed differentially accessible regions (DAR) within each cluster. We observed decreased accessibility in promoter, 5’ UTR, downstream, and intronic regions in HSC and CD14+ clusters with CHD (**Supp. Figure 5F**). Functional enrichment using Enricher and Metascape of promoters, 5’ UTR, and downstream regions that were more open in HSC in control animals (closed in CHD group) harbored genes important for “regulation of IL10 production”, “mononuclear cell differentiation” and “positive regulation of NIK/NF-kappaB signaling”, while those more open with CHD played a role in “regulation of T-helper 1 cell cytokine production”, “regulation of type I interferon-mediated signaling pathway”, and “regulation of defense response” indicating increased inflammation with CHD (**Figure 5D-E**). Functional enrichment of these same regions in CD14+ cells from control animals mapped to terms such as “negative regulation of leukocyte apoptotic process”, “dendritic cell differentiation”, “regulation of MAP kinase activity”, and “regulation of interleukin 12 production”, while those from CHD animals mapped to apoptotic processes as well as “regulation of protein localization to the nucleolus” and “interleukin 6 mediated signaling pathways” (**Figure 5F-G**). GREAT analysis of DARs within distal intergenic regions that were open in CD14+ cells from control animals showed enrichment to “positive regulation of leukocyte cell−cell adhesion”, “negative regulation of MAPK cascade”, and “IFN−gamma−mediated signaling pathway” while those more open in CHD cells mapped to “negative regulation of catabolic process” (**Figure 5F**). These observations indicate that CHD impacts monocytes and their progenitors in the bone marrow on an epigenetic level, which could play a role in the altered differentiation trajectory of these cells and lead to observed functional complications.

## DISCUSSION

Chronic consumption of alcohol can disrupt anti-microbial defenses and exacerbate inflammatory responses of monocytes and macrophages (14, 16, 17, 36). Using a NHP model of voluntary ethanol consumption, we found increased percentages of blood monocytes and splenic macrophages relative to total cells with CHD compared to control animals (14, 36). In addition to increased monocyte percentages in the periphery, we and others have noted an increase in inflammatory monocyte responses to stimulation with CHD (14, 16, 17, 36). As monocytes are short-lived cells in the periphery, these observations suggest that chronic drinking could lead to dysregulation of monocyte maturation and/or export from the bone marrow. Other studies have suggested that ethanol and its metabolites modulate monocyte function through epigenetic modification and oxidative stress of both mature cells and hematopoietic progenitors, but the exact mechanisms remain poorly defined (14, 34, 36). Therefore, in this study, we assess the phenotypic, functional, and single cell transcriptomic and epigenomic profiles of bone-marrow resident monocytes/macrophages and CD34+ HSC collected from NHP after 12 months of voluntary CHD.

While we noted no differences in percentages of mature monocytes by flow cytometry in the bone marrow with CHD, scRNA-Seq showed increased expression of inflammatory genes across al CD14+ bone-marrow resident cells subsets. Interestingly, the frequency of a population of *CXCR4*hi monocytes, which may be transitional pre-monocytes that help to maintain the mature monocyte pool in the bone marrow (38), was increased with CHD. The expression of *CCR2,* which plays a critical role in monocyte egress from the bone marrow (39), was also increased on bone marrow resident classical monocytes with CHD. As CCR2 is critical for monocyte egress from the bone marrow, this observation suggests CHD-induced disruption of bone marrow monocyte trafficking between the bone marrow and blood (39).

Prior studies have suggested alterations in progenitor populations with CHD, but none to our knowledge have examined monopoiesis in this context. We performed *in vitro* maturation assays of CD34+ HSC and found skewing towards the GMP (neutrophil-monocyte) lineage and away from MDP (DC-monocyte) lineage. This skewing was accompanied by a significant increase in inflammatory mediator production from these culture-derived monocytes. A second assay also showed increased production of colonies containing granulocyte/monocyte progenitor cells with CHD consumption. These data suggest CHD-induced shifts in myelopoiesis and monopoiesis from CD34+ progenitors, giving rise to monocytes poised towards a heightened inflammatory state.

CHD metabolism produces acetaldehyde and increases NADH levels, which in turn promotes reactive oxygen species (ROS) production (43). These products induce DNA damage as well as oxidative stress on cells and can have direct effects on cellular function (43, 44). Indeed, differential expression analysis from scRNA-Seq data revealed increases in transcriptional signatures of oxidative stress, hypoxia, and chronic inflammation in bone marrow resident CD14+ and CD34+ cells. Additionally, expression of several mitochondrial genes were increased in monocyte progenitors, suggesting CHD-induced shifts to cellular metabolism in the bone marrow. These observations are in line with an earlier study in NHP bone marrow reported altered mitochondrial function with daily CHD (34) and a recent study reporting increased mitochondrial in alveolar macrophages with CHD (45). Collectively, these data suggest that the hyper-inflammatory profiles of circulating monocytes with CHD could be attributed to CHD-induced oxidative stress on CD34+ progenitors in the bone marrow.

Previous work in peripheral monocytes and tissue-resident macrophages showed significant epigenetic changes associated with CHD (14, 36, 45). In this study, we identified few differentially accessible regions with CHD in bone marrow resident cells. One potential explanation is that the depth of the scATAC-Seq analysis was not sufficient to identify DAR within smaller clusters. Alternatively, the epigenetic changes detected in the periphery may be primarily driven by ethanol and its metabolites which would be more concentrated in circulation. We did observe some altered DAR in cis-regulatory regions as well as an increase in terms associated with inflammation in HSC and CD14+ cells. Most notably, accessibility of regions overlapping genes that regulate or respond to IL-6 was increased.

One of the mechanisms by which CHD could lead to HSCP and bone marrow niche dysfunction is through altering the bone marrow CD169+ macrophage population, which are critically important for maintenance of the niche (40, 41). Expression of activation marker CD86 was decreased in macrophages with CHD at resting state and upregulation of HLA-DR after bacterial agonist stimulation was dampened. This was accompanied by heightened transcriptional signatures of oxidative phosphorylation. Together with previous reports of reduced macrophage functional capacities with alcohol exposure including phagocytosis (46), these data suggest that the ability of these macrophages to maintain the bone marrow niche are compromised with CHD.

The data from this study strongly suggests that CHD alters monopoiesis from the bone marrow compartment in NHPs. Similar to circulatory monocytes, CD14+ cells that reside in the bone marrow compartment have inflammatory-skewed transcriptional and functional profiles, suggesting CHD remodeling of this tissue compartment. Mechanistically, CHD disrupts the differentiation of CD34+ cells into monocytes leading to the *in vitro* production of monocytes with a hyper-inflammatory phenotype. These disruptions are potentially mediated by CHD-induced oxidative stress shown transcriptionally and through altered profiles of niche-maintaining macrophages. Future research efforts will focus on determining a timeline for alcohol-induced bone marrow remodeling and determining how alcohol alters monocyte release from the bone marrow compartment.

## METHODS AND MATERIALS

### Animal studies and sample collection

These studies used samples from an NHP model of ethanol self-administration established through schedule-induced polydipsia (47–49). Briefly, in this model, rhesus macaques are introduced to a 4% w/v ethanol solution during a 90-day induction period followed by concurrent access to the 4% w/v solution and water for 22 hours/day for one year. During this time, the macaques adopt a stable drinking phenotype defined by the amount of ethanol consumed per day and the pattern of ethanol consumption (g/kg/day) (47). Blood samples were taken from the saphenous vein every 5-7 days at 7 hrs after the onset of the 22 hrs/day access to ethanol and assayed by headspace gas chromatography for blood ethanol concentrations (BECs).

For these studies, blood and bone marrow samples were collected through the Monkey Alcohol Tissue Research Resource (www.matrr.com) where more information about the animals can be obtained. Blood samples (2 timepoints) were collected from 7 male rhesus macaques with 3 animals serving as controls and 4 classified as heavy drinkers based on 12-month daily averages of ethanol self-administration (Cohort 14 on www.matrr.com). Bone marrow samples were collected from one male cohort of 8 animals and one female cohort of 9 animals for a total of 7 controls and 10 heavy drinkers (Cohorts 6a and 7a on www.matrr.com). Data from Cohorts 6 and 7a have been reported in three previous studies of innate immune system response to alcohol (14–16). Peripheral Blood Mononuclear Cells (PBMC) bone marrow cells were cryopreserved until they could be analyzed as a batch. The average daily ethanol intake and other cohort information for each animal is outlined in **Supp. Table 1**.

### Flow cytometry analysis

0.5-1×10^6^ PBMC were stained with the following surface antibodies against: CD14 (Biolegend, M5E2, AF700), HLA-DR (Biolegend, L243, APC-Cy7) to identify monocytes. 0.5-1×10^6^ bone marrow cells were stained with the following surface antibodies (2 panels) against: CD14 (Biolegend, M5E2, AF700), HLA-DR (Biolegend, L243, APC-Cy7), CD16 (Biolegend, 3G8, PB), CD80 (Biolegend, 2D10, PE), CD86 (Biolegend, IT2.2, BV605), CD169 (Biolegend, 7-239, PE-Cy7), and Sytox Green to identify monocyte and macrophage populations; CD20 (Biolegend, 2H7, BV510), CD3 (BD Biosciences, SP34-2, V500), CD14 (Biolegend, M5E2, AF700), CD38 (StemCell, AT-1, FITC), CD123 (Biolegend, 6H6, PerCP-Cy5.5), CD34 (Biolegend, 561, PE-Cy7), CD45RA (Miltenyi, T6D11, PE), CD64 (Biolegend, 10.1, BV711), CD90 (Biolegend, 5E10, APC), CD127 (Miltenyi, PE-Vio615), and Sytox Blue to identify bone marrow progenitor populations. Progenitor populations were defined as: HSC (Lin.-CD34+CD38-CD45RA-CD90+), MPP(Lin.-CD34+CD38-CD45RA-CD90-), CLP(Lin.-CD34+CD38+CD127+), CMP(Lin.-CD34+CD38+CD45RA-CD123+), MEP (Lin.-CD34+CD38+CD45RA-CD123-), GMP(Lin.-CD34+CD38+CD45RA+CD123+CD64-), cMoP(Lin.-CD34+CD38+CD45RA+CD123+CD64+). All samples were acquired with an Attune NxT Flow Cytometer (ThermoFisher Scientific, Waltham, MA) and analyzed using FlowJo software (Ashland, OR). Percentage of live cells or Median Fluorescence Intensities (MFI) were assessed for each marker.

### PBMC and bone marrow Stimulation Assays

1×10^6^ freshly thawed PBMC were cultured in RPMI supplemented with 10% FBS with or without 1 ug/mL LPS (TLR4 ligand, *E.coli* 055:B5; Invivogen, San Diego CA) and Brefeldin A for 6 hours in 96-well tissue culture plates at 37C in a 5% CO_2_ environment. They were next stained with an antibody cocktail of CD20 (Biolegend, 2H7, BV510), CD3 (BD Biosciences, SP34-2, V500), CD14 (Biolegend, M5E2, AF700), and HLA-DR (Biolegend, L243, APC-Cy7) and Fixable Yellow Live/Dead stain. Stained cells were then fixed and permeabilized using Fixation buffer (BioLegend) and incubated overnight with intracellular antibody TNF (BD Biosciences, MAb11, APC).

1×10^6^ freshly thawed bone marrow cells were cultured in RPMI supplemented with 10% FBS with or without a bacterial agonist cocktail (2ug/mL Pam3CSK4 (TLR1/2 agonist, InvivoGen), 1 ug/mL FSL-1 (TLR2/6 agonist, Sigma Aldrich), and 1 ug/mL LPS (TLR4 agonist from E. coli O111:B4, InvivoGen)) and Brefeldin A for 6 hours in 96-well tissue culture plates at 37C in a 5% CO_2_ environment. They were next stained with an antibody cocktail of CD14 (Biolegend, M5E2, AF700), HLA-DR (Biolegend, L243, APC-Cy7), CD34 (Biolegend, 561, PE-Cy7), CD16 (Biolegend, 3G8, PB), and CD169 (Biolegend, 7-239. Stained cells were then fixed and permeabilized using Fixation buffer (BioLegend) and incubated overnight with a cocktail of intracellular antibodies – IL-6 (BD Biosciences, MQ2-6A3, FITC), TNFα (BD Biosciences, MAb11, APC), CCL4 (MIP-1β) (BD Biosciences, D21-1351, PE).

All samples were acquired with an Attune NxT Flow Cytometer (ThermoFisher Scientific, Waltham, MA) and analyzed using FlowJo software (Ashland, OR).

### Monocyte Differentiation Assay

1×10^3^ sorted CD34+ cells/well were plated in a 96-well plate in 100uL StemSpan SFEM Media supplemented with StemSpan Myeloid Expansion Supplement II containing TPO, SCF, Flt3, GM-CSF, M-CSF and incubated at 37C in a 5% CO_2_ environment. On culture day 4, the volume of the cultures was brought up to 200uL with the same media. On culture day 7, the cultures were incubated with or without a bacterial agonist cocktail (2ug/mL Pam3CSK4 (TLR1/2 agonist, InvivoGen), 1 ug/mL FSL-1 (TLR2/6 agonist, Sigma Aldrich), and 1 ug/mL LPS (TLR4 agonist from E. coli O111:B4, InvivoGen)) for 6 hours. Supernatants were collected and stored short-term at -80C. The cells were further stained with an antibody cocktail of: CD34 (Biolegend, 561, PE-Cy7), CD14 (Biolegend, M5E2, AF700), HLA-DR (Biolegend, L243, APC-Cy7), CD11C (Invitrogen, 3.9, PE-eFluor610), CD115 (Biolegend, 9-4D2-1E4, PE), and CCR2 (R&D, 48607, PerCP-Cy5.5). All samples were acquired with an Attune NxT Flow Cytometer (ThermoFisher Scientific, Waltham, MA) and analyzed using FlowJo software (Ashland, OR).

### Luminex Assay

Immune mediators in the supernatants from monocyte cultures were measured using a Milliplex Multiplex assay panel measuring levels of TNFα, IL-6, IL-1β, G-CSF, CCL3(MIP-1α), and IL-2 (Millipore, Burlington, MA). Differences in induction of proteins post stimulation were calculated using log_10_(pg/mL fold-change +1). Undiluted samples run in duplicates on the Magpix Instrument (Luminex, Austin, TX). Data were fit using a 5P-logistic regression on xPONENT software (version 7.0c).

### Colony Forming Unit Assay

MethoCult colony-forming unit (CFU) assay was performed with MethoCult H4435-enriched medium (STEMCELL Technologies, Vancouver, Canada) following manufacturer’s protocol. Briefly, CD34+ cells were FACS sorted from freshly thawed bone marrow cells and resuspended in IMDM+2% FBS to achieve a final plating concentration of 1000 cells per culture dish. Cells were gently mixed with 3mL MethoCult and plated into 35mm culture dishes in triplicate. Cultures were incubated at 37C in a 5% CO_2_ environment. Total colonies were counted on days 7 and 10, and additional colony identification was performed on day 10.

### Single cell RNA library preparation and sequencing – CD14+

Bone marrow cells from control females (n=3 pooled), control males (n=4), CHD females (n=3 pooled), and CHD males (n=4), were thawed and subjected to the following downstream protocols. CD14+ cells (mature monocytes) were sorted using a BD FACSAria Fusion. Sorted CD14+ cells from the control and CHD groups of female animals were pooled and resuspended at a concentration of 1,200 cells/uL and loaded into the 10X Chromium Controller aiming for an estimated 10,000 cells per sample. cDNA amplification and library preparation (10X v3 chemistry) were performed for samples according to manufacturer protocol and sequenced on a NovaSeq S4 (Illumina) to a depth of >30,000 reads/cell.

Freshly thawed bone marrow cells from male animals were first incubated with TruStain FcX for 10 minutes followed by surface staining with anti-CD14 antibody. Cells were washed in PBS + 0.04% BSA and then incubated with 10X CellPlex oligos (10X Genomics) for 5 minutes. CD14+ cells were sorted into respective control or CHD pools for live, CD14+ cells using a BD FACSAria Fusion. After sorting, pools were resuspended at a concentration of 1,600 cells/uL and loaded into the 10X Chromium Controller aiming for an estimated 18,000 cells per sample. cDNA amplification and library preparation (10X v3.1 dual index chemistry) were performed for samples according to manufacturer protocol and sequenced on a NovaSeq S4 (Illumina) to a depth of >30,000 reads/cell. Cell multiplexing libraries were constructed according to manufacturer protocol and sequenced on a NovaSeq S4 (Illumina) to a depth of >5,000 reads/cell.

### Single cell RNA library preparation-monocyte differentiation cultures

Monocyte differentiation culture cells from control (n=2 male + 2 female pooled) and CHD (n=2 male + 2 female pooled) cultures were counted for viability (>70%) and pooled according to group. Pools were resuspended at a concentration of 1,200 cells/uL and loaded into the 10X Chromium Controller aiming for an estimated 10,000 cells per sample. cDNA amplification and library preparation (10X v3 chemistry) were performed for samples according to manufacturer protocol and sequenced on a NovaSeq S4 (Illumina) to a depth of >30,000 reads/cell.

### Single cell RNA library preparation and sequencing-CD34+

Bone marrow cells from control females (n=3), control males (n=4), CHD females (n=3), and CHD males (n=4), were thawed and stained with anti-CD34 antibodies and sorted for live CD34+ cells on a BD FACSAria Fusion. Sorted CD34+ samples from female animals were resuspended at a concentration of 1,200 cells/uL and loaded into the 10X Chromium Controller aiming for an estimated 10,000 cells per sample. cDNA amplification and library preparation (10X v3 chemistry) were performed for samples according to manufacturer protocol and sequenced on a NovaSeq S4 (Illumina) to a depth of >30,000 reads/cell.

Freshly thawed bone marrow cells from male animals were first incubated with TruStain FcX for 10 minutes followed by surface staining with anti-CD34 antibody. Cells were washed in PBS + 0.04% BSA and then incubated with 10X CellPlex oligos (10X Genomics) for 5 minutes. CD34+ cells were sorted into respective control or CHD pools for live, CD34+ cells on a BD FACSAria Fusion. After sorting, pools were resuspended at a concentration of 1,600 cells/uL and loaded into the 10X Chromium Controller aiming for an estimated 18,000 cells per sample. cDNA amplification and library preparation (10X v3.1 dual index chemistry) were performed for samples according to manufacturer protocol and sequenced on a NovaSeq S4 (Illumina) to a depth of >30,000 reads/cell. Cell multiplexing libraries were constructed according to manufacturer protocol and sequenced on a NovaSeq S4 (Illumina) to a depth of >5,000 reads/cell.

### Single cell RNA-Seq data analysis

Sequencing reads were aligned to the Mmul_8.0.1 reference genome using cellranger v6.0.1 (50) (10X Genomics) using the *count* function for single sample libraries and the *multi* function for multiplexed samples. Quality control steps were performed prior to downstream analysis with *Seurat* (51), filtering out cells with fewer than 200 unique features (ambient RNA) and cells with greater than 20% mitochondrial content (dying cells). Data normalization was performed using *SCTransform* (52), correcting for differential effects of mitochondrial and cell cycle gene (only CD14+ dataset) expression levels. Sample integration was performed using the *SelectIntegrationFeatures* (using 3000 features), *PrepSCTIntegration*, *FindIntegrationAnchors*, and *IntegrateData* functions. Clustering was performed using the first 20, 10, and 30 principal components for the CD14+, monocyte differentiation, and CD34+ datasets, respectively. Contaminating clusters with an over-representation of B or T cell gene expression were removed for downstream analysis. Clusters were characterized into distinct subsets using the *FindAllMarkers* function (**Supp. Table 2**). Figures were generated using *Seurat, ggplot2*, and *pheatmap*.

### Differential expression analyses

Differential expression analysis (CHD relative to Control) was performed using MAST under default settings in *Seurat*. Only statistically significant genes (Log_10_(Fold change) cutoff ≥ 0.25; adjusted p-value≤ 0.05) were included in downstream analysis.

### Pseudo-temporal analysis

Pseudotime trajectory of the monocyte differentiation and CD34+ datasets were reconstructed using *Slingshot* (53). The UMAP dimensional reduction performed in *Seurat* was used as the input for *Slingshot*. For calculation of the lineages and pseudotime, the progenitor/HSC population was selected as the start. Temporally expressed genes were identified by ranking all genes by their variance across pseudotime and then further fit using the generalized additive model (GAM) with pseudotime as an independent variable.

### Module Scoring and functional enrichment

For gene scoring analysis, we compared gene signatures and pathways from KEGG (https://www.genome.jp/kegg/pathway.html) and the Human Cell Atlas bone marrow single cell analysis (42) (**Supp. Table 3**) in the clusters/groups using *Seurat’s AddModuleScore* function. Values for module scores were further exported from *Seurat* and tested for significance in Prism 7. Over representative gene ontologies were identified by enrichment of differential signatures using Metascape. All plots were generated using *ggplot2* and *Seurat*.

### RNA isolation and library preparation for bulk RNA-Seq

Total RNA was isolated from sorted CMP/GMP using the mRNeasy kit (Qiagen, Valencia CA) following manufacturer instructions and quality assessed using Agilent 2100 Bioanalyzer. Libraries from purified progenitor RNA were generated using the NEBnext Ultra II Directional RNA Library Prep Kit for Illumina (NEB, Ipswitch, MA, USA). rRNA depleted RNA was fragmented, converted to double-stranded cDNA and ligated to adapters. The roughly 300bp-long fragments were then amplified by PCR and selected by size exclusion. Libraries were multiplexed and following quality control for size, quality, and concentrations, were sequenced to an average depth of 20 million 100bp reads on the NovaSeq S4 platform.

### Bulk RNA-Seq data analysis

RNA-Seq reads were quality checked using FastQC (https://www.bioinformatics.babraham.ac.uk/projects/fastqc/), adapter and quality trimmed using TrimGalore(https://www.bioinformatics.babraham.ac.uk/projects/trim_galore/), retaining reads at least 35bp long. Reads were aligned to *Macaca mulatta* genome (Mmul_8.0.1) based on annotations available on ENSEMBL (Mmul_8.0.1.92) using TopHat (54) internally running Bowtie2 (55). Aligned reads were counted gene-wise using GenomicRanges (56), counting reads in a strand-specific manner. Genes with low read counts (average <5) and non-protein coding genes were filtered out before differential gene expression analyses. Raw counts were used to test for differentially expressed genes (DEG) using edgeR (57), defining DEG as ones with at least two-fold up or down regulation and an FDR controlled at 5%. edgeR analysis is provided in **Supp. Table 3.4**.

### Single cell ATAC-Seq library preparation and sequencing

Bone marrow cells from control males (n=3), CHD females (n=2), and CHD males (n=2), were thawed and stained with anti-CD34 and anti-CD14 antibodies and Sytox Green to sort for live CD34+ cells and live CD14+ cells on a BD FACSAria Fusion. Sorted cells for each sample were lysed, nuclei isolated, nuclei counted, and transposition reaction performed according to manufacturer’s instructions (10X Genomics). 3,000-20,000 nuclei were obtained per sample and loaded into the 10X Chromium Controller. Library preparation (10X v2 chemistry) was performed for samples according to manufacturer protocol and sequenced on a NovaSeq S4 (Illumina) to a depth of >25,000 paired reads/cell.

### Single cell ATAC-Seq data processing, dimensional reduction, and clustering

Sequencing reads were pre-processed using the cellranger-atac pipeline (v2.1.0) (10X Genomics) where accessibility counts for each cell were aligned to the Mmul_8.0.1 reference genome. The ArchR package (58) (v1.0.1) was used for downstream analysis following their vignette in R (v4.1.0). Arrow files were created from each sample fragment file and low-quality cells were filtered out (<3162 fragments, <8 TSS enrichment, doublets calculated by addDoubletScores). An ArchR project was created by combining all Arrow files. Iterative latent semantic indexing (LSI) was performed as the first dimensional reduction followed by the *addHarmony* function to correct batch effects. Uniform manifold approximation and projection (UMAP) was used for the final dimensional reduction and clusters were added using the *addClusters* function. Marker features for each cluster based on gene scores were identified using the *getMarkerFeatures* function. Myeloid cell lineage clusters were defined by accessible marker genes, subset, and the above dimensional reduction and clustering steps were performed again. Pseudo-bulk replicates were created for peak calling using MACS2. Per-cell deviations across motif annotations were computed using the *addDeviationsMatrix* function. The *getMarkerFeatures* function was used to determine differentially accessible regions (DAR) between CHD and Control clusters or groups of clusters. All UMAP and heatmap plots were generated using ArchR functions.

### Single cell ATAC-Seq trajectory analysis

Trajectory analysis was performed on HSC, GMP, MP/GMP, MP, and monocyte clusters from the control and CHD groups using the *addTrajectory* function in ArchR. To identify positive TF regulators of the trajectory, we performed an integrative analysis of gene scores and motif accessibility across the identified psuedotime trajectory.

### Promoter and Distal Intergenic Peak Annotation and Functional Enrichment Analysis

Genomic annotation of open chromatin regions related to the promoter, 5’ UTR, downstream, and distal intergenic regions in DAR analysis was assigned using ChIPSeeker. Promoters were defined using the following criteria: −1000 bp to +100 bp around the transcriptional start site (TSS). Genes with no annotations were excluded from downstream analyses. Functional enrichment analysis was performed for the promoter, 5’ UTR, and downstream regions using Metascape (https://metascape.org) and Enrichr (https://maayanlab.cloud/Enrichr/). The liftOver strategy was carried out to convert the distal intergenic regions of the macaque to the human genome (hg38) coordinates using UCSC liftOver tool due to the lack of available macaque annotation databases. This strategy has been recently reported as an alternative for nonhuman primates’ studies because instead of qualitative differences, quantitative differences in enhancer activity are the prevalent source of regulatory landscape divergence among closely related species (59). The functions of cis-regulatory regions were predicted using GREAT (http://great.stanford.edu/public/html/).

### Statistical Analysis

All statistical analyses were conducted in Prism 7(GraphPad). Data sets were first tested for normality and outliers. Two group comparisons were carried out an unpaired t-test with Welch’s correction or Mann-Whitney test. Differences between 3 groups were tested using one-way ANOVA (α=0.05) followed by Holm Sidak’s multiple comparisons tests. Error bars for all graphs are defined as ± SEM. Statistical significance of functional enrichment was defined using hypergeometric tests. P-values less than or equal to 0.05 were considered statistically significant. Values between 0.05 and 0.1 are reported as trending patterns.

### Study Approval

This study was approved by the Oregon National Primate Research Center (ONPRC) Institutional Animal Care and Use Committee (IACUC). ONPRC is an Association for Assessment and Accreditation of Laboratory Animal Care (AAALAC) approved institute. The work described in this study has been approved by the Institutional Biosafety Committee (IBC) of all associated institutions.

## Author Contributions

S.A.L., K.A.G., and I.M. conceived and designed the experiments. S.A.L. and B.M.D. performed the experiments. S.A.L., B.M.D., M.B, and Q.Q analyzed the data. S.A.L. and I.M. wrote the paper. All authors have read and approved the final draft of the manuscript.

## Acknowledgements

We are grateful to the members of the Grant laboratory for expert animal care and sample procurement. We thank Dr. Jennifer Atwood for assistance with sorting in the flow cytometry core at the Institute for Immunology, UCI. We thank Dr. Melanie Oakes from UCI Genomics and High-Throughput Facility for assistance with sequencing. We thank Dr. Helen Goodridge for assistance with CFU assay protocols and analysis.

## Funding

This study was supported by NIH 1R01AA028735-01 (Messaoudi), 5U01AA013510-20 (Grant), and 2R24AA019431-11 (Grant). S.A.L is supported by NIH 1F31A028704-01. The content is solely the responsibility of the authors and does not necessarily represent the official views of the NIH.

## Competing interests

No competing interests reported.

## Data availability

The datasets supporting the conclusions of this article are available on NCBI’s Sequence Read Archive (SRA# pending).

## FIGURE LEGENDS

**Supp. Figure 1: scRNA-Seq of CD14+ cells from the bone marrow**

A) Example gating strategy for monocyte populations from the bone marrow. B) UMAPs of each individual macaque sample. C) UMAP annotated by broad classification of cell type and corresponding pie charts of abundance of cells from each group. D) Stacked violin plots of genes related to GMP or MDP lineages across each classical cluster and split by CHD and control groups. Statistical analysis performed by Mann-Whitney test where *=p<0.05, **=p<0.01, ***=p<0.001, ****=p<0.0001. E) Bar plot representing -log_10_(q-value) functional enrichment scores for genes up-(blue, top) and downregulated (green, bottom) in non-classical monocytes with CHD. F,G) Violin plots of down-(F) and up-(G) regulated DEG with CHD in the macrophage cluster.

**Supp. Figure 2: CD34+ progenitor differentiation assays and flow cytometry of CD34+ cells from the bone marrow**

A) Bar plots showing percentage of CD115+ (left) and CD11C+ (right) in the indicated culture populations from flow cytometry. B) Bar graphs showing average colony numbers on day 7 and day 10 of culture. C) Gating strategy for bone marrow progenitor cells. D) Bar plots showing percentages of progenitor populations in the bone marrow compartment determined by flow cytometry. Unless indicated, statistical significance was tested by t-test with Welch’s correction. *=p<0.05, **=p<0.01, ****=p<0.0001.

**Supp. Figure 3: scRNA-Seq of CD34+ cells from the bone marrow**

A) UMAP clustering of 41,125 cells and indicated cluster identification. B) Heatmap showing averaged gene expression of highly expressed genes from each progenitor cluster. C) Stacked violin plot showing expression of genes identified using Seurat’s *FindAllMarkers* function. D) Feature plots showing relative expression of module scores for CMP, Granulocyte progenitors, MDP-1, and MDP-1 in the myeloid lineage. E) Box plots of percentages of each cluster across control and CHD groups. Statistical analysis performed by two-way ANOVA with multiple comparisons, *=p<0.05.

**Supp. Figure 4: Myeloid progenitor subsets in the bone marrow**

A) UMAP projection of integrated CD34+ (Figure 4) and CD14+ (Figure 2) datasets (left). Feature plot of *CD14* expression to identify contaminating CD14+ cells (middle). UMAP of Seurat clusters labeled with CD14 or CD34 designation for selection of only CD34+ cells (right). Heatmaps of genes explaining the DC (left) and neutrophil (right) lineage trajectories. C) Cell density plots for Control and CHD groups across the DC (top) and neutrophil (bottom) trajectory lineages determined by Slingshot. D,F) Bar plot showing functional enrichment terms from less (D) and more (F) mature clusters. E,G) Heatmap showing averaged gene expression of upregulated DEG from less (E) and more (G) mature clusters split by CHD and control groups.

**Supp. Figure 5: scATAC-Seq of bone marrow CD34+ and CD14+ cells**

CD34+ and CD14+ cells were sorted from bone marrow and subjected to 10X scATAC-Seq. A) UMAP projection of all cells based on gene accessibility. B) Feature plots of indicated marker genes from the gene score matrix. C) Stacked bar plot distribution of clusters across each sample. D) UMAP plots of motif accessibility for each of the indicated transcription factors. E) Bar plot of percentages of control and CHD groups across each cluster. F) Bar plots of the percent of differentially accessible regions (left) and total numbers of differentially accessible regions (right) between control and CHD samples.

**Supp. Table 1: Animal cohort characteristics**

**Supp. Table 2: FindAllMarkers (CD14+, Culture, CD34+)**

**Supp. Table 3: Module Score Gene Lists**

**Supp. Table 4: Marker GeneScores scATACseq**

## Notes

### Competing Interest Statement

The authors have declared no competing interest.

## REFERENCES

1. (WHO) WHO. 2018.

2. O’Keefe JH, et al. Alcohol and cardiovascular health: the dose makes the poison…or the remedy. Mayo Clin Proc. 2014;89(3):382–93.

3. Mukamal KJ, and Rimm EB. Alcohol’s effects on the risk for coronary heart disease. Alcohol Res Health. 2001;25(4):255–61.

4. Fedirko V, et al. Alcohol drinking and colorectal cancer risk: an overall and dose-response meta-analysis of published studies. Ann Oncol. 2011;22(9):1958–72.

5. Baan R, et al. Carcinogenicity of alcoholic beverages. Lancet Oncol. 2007;8(4):292–3.

6. Grewal P, and Viswanathen VA. Liver cancer and alcohol. Clin Liver Dis. 2012;16(4):839–50. 7.

7. Priddy BM, et al. Sex, strain, and estrous cycle influences on alcohol drinking in rats. Pharmacol Biochem Behav. 2017;152:61–7.

8. Bruha R, et al. Alcoholic liver disease. Prague Med Rep. 2009;110(3):181–90.

9. O’Brien JM, Jr., et al. Insurance type and sepsis-associated hospitalizations and sepsis-associated mortality among US adults: a retrospective cohort study. Crit Care. 2011;15(3):R130.

10. Delgado-Rodriguez M, et al. Alcohol consumption and the risk of nosocomial infection in general surgery. Br J Surg. 2003;90(10):1287–93.

11. Saitz R, et al. The impact of alcohol-related diagnoses on pneumonia outcomes. Arch Intern Med. 1997;157(13):1446–52.

12. Baum MK, et al. Alcohol use accelerates HIV disease progression. AIDS Res Hum Retroviruses. 2010;26(5):511–8.

13. Bhattacharya R, and Shuhart MC. Hepatitis C and alcohol: interactions, outcomes, and implications. J Clin Gastroenterol. 2003;36(3):242–52.

14. Lewis SA, et al. Transcriptional, Epigenetic, and Functional Reprogramming of Monocytes From Non-Human Primates Following Chronic Alcohol Drinking. Front Immunol. 2021;12:724015.

15. Sureshchandra S, et al. Transcriptome Profiling Reveals Disruption of Innate Immunity in Chronic Heavy Ethanol Consuming Female Rhesus Macaques. PLoS One. 2016;11(7):e0159295.

16. Sureshchandra S, et al. Dose-dependent effects of chronic alcohol drinking on peripheral immune responses. Sci Rep. 2019;9(1):7847.

17. Szabo G, and Saha B. Alcohol’s Effect on Host Defense. Alcohol Res. 2015;37(2):159–70.

18. Kawamura S, and Ohteki T. Monopoiesis in humans and mice. Int Immunol. 2018;30(11):503–9.

19. Wolf AA, et al. The Ontogeny of Monocyte Subsets. Front Immunol. 2019;10:1642.

20. Yanez A, et al. Granulocyte-Monocyte Progenitors and Monocyte-Dendritic Cell Progenitors Independently Produce Functionally Distinct Monocytes. Immunity. 2017;47(5):890–902 e4.

21. Kawamura S, et al. Identification of a Human Clonogenic Progenitor with Strict Monocyte Differentiation Potential: A Counterpart of Mouse cMoPs. Immunity. 2017;46(5):835–48 e4.

22. Lee J, et al. Restricted dendritic cell and monocyte progenitors in human cord blood and bone marrow. J Exp Med. 2015;212(3):385–99.

23. Teh YC, et al. Capturing the Fantastic Voyage of Monocytes Through Time and Space. Front Immunol. 2019;10:834.

24. Takizawa H, et al. Demand-adapted regulation of early hematopoiesis in infection and inflammation. Blood. 2012;119(13):2991–3002.

25. Baldridge MT, et al. Inflammatory signals regulate hematopoietic stem cells. Trends Immunol. 2011;32(2):57–65.

26. Shi X, et al. Alcohol abuse and disorder of granulopoiesis. Pharmacol Ther. 2019;198:206–19.

27. Panasiuk A, and Kemona A. Bone marrow failure and hematological abnormalities in alcoholic liver cirrhosis. Rocz Akad Med Bialymst. 2001;46:100–5.

28. Liu YK. Effects of alcohol on granulocytes and lymphocytes. Semin Hematol. 1980;17(2):130–6.

29. Latvala J, et al. Excess alcohol consumption is common in patients with cytopenia: studies in blood and bone marrow cells. Alcohol Clin Exp Res. 2004;28(4):619–24.

30. Ballard HS. The hematological complications of alcoholism. Alcohol Health Res World. 1997;21(1):42–52.

31. Smith C, et al. The effects of alcohol and aldehyde dehydrogenases on disorders of hematopoiesis. Adv Exp Med Biol. 2015;815:349–59.

32. Raasch CE, et al. Acute alcohol intoxication impairs the hematopoietic precursor cell response to pneumococcal pneumonia. Alcohol Clin Exp Res. 2010;34(12):2035–43.

33. Shi X, et al. Impairment of Hematopoietic Precursor Cell Activation during the Granulopoietic Response to Bacteremia in Mice with Chronic-Plus-Binge Alcohol Administration. Infect Immun. 2017;85(11).

34. Varlamov O, et al. Daily Ethanol Drinking Followed by an Abstinence Period Impairs Bone Marrow Niche and Mitochondrial Function of Hematopoietic Stem/Progenitor Cells in Rhesus Macaques. Alcohol Clin Exp Res. 2020;44(5):1088–98.

35. Siggins RW, et al. Dysregulation of myelopoiesis by chronic alcohol administration during early SIV infection of rhesus macaques. Alcohol Clin Exp Res. 2014;38(7):1993–2000.

36. Sureshchandra S, et al. Chronic heavy drinking drives distinct transcriptional and epigenetic changes in splenic macrophages. EBioMedicine. 2019;43:594–606.

37. Allen DC, et al. Effect of repeated abstinence on chronic ethanol self-administration in the rhesus monkey. Psychopharmacology (Berl). 2018;235(1):109–20.

38. Chong SZ, et al. CXCR4 identifies transitional bone marrow premonocytes that replenish the mature monocyte pool for peripheral responses. J Exp Med. 2016;213(11):2293–314.

39. Tsou CL, et al. Critical roles for CCR2 and MCP-3 in monocyte mobilization from bone marrow and recruitment to inflammatory sites. J Clin Invest. 2007;117(4):902–9.

40. Winkler IG, et al. Bone marrow macrophages maintain hematopoietic stem cell (HSC) niches and their depletion mobilizes HSCs. Blood. 2010;116(23):4815–28.

41. Mitroulis I, et al. Regulation of the Bone Marrow Niche by Inflammation. Front Immunol. 2020;11:1540.

42. Hay SB, et al. The Human Cell Atlas bone marrow single-cell interactive web portal. Exp Hematol. 2018;68:51–61.

43. Di Rocco G, et al. Stem cells under the influence of alcohol: effects of ethanol consumption on stem/progenitor cells. Cell Mol Life Sci. 2019;76(2):231–44.

44. Garaycoechea JI, et al. Genotoxic consequences of endogenous aldehydes on mouse haematopoietic stem cell function. Nature. 2012;489(7417):571–5.

45. Lewis SA, et al. Ethanol Consumption Induces Non-Specific Inflammation and Functional Defects in Alveolar Macrophages. Am J Respir Cell Mol Biol. 2022.

46. Karavitis J, and Kovacs EJ. Macrophage phagocytosis: effects of environmental pollutants, alcohol, cigarette smoke, and other external factors. J Leukoc Biol. 2011;90(6):1065–78.

47. Baker EJ, et al. Chronic alcohol self-administration in monkeys shows long-term quantity/frequency categorical stability. Alcohol Clin Exp Res. 2014;38(11):2835–43.

48. Grant KA, et al. Drinking typography established by scheduled induction predicts chronic heavy drinking in a monkey model of ethanol self-administration. Alcohol Clin Exp Res. 2008;32(10):1824–38.

49. Jimenez VA, et al. An ultrastructural analysis of the effects of ethanol self-administration on the hypothalamic paraventricular nucleus in rhesus macaques. Front Cell Neurosci. 2015;9:260.

50. Zheng GX, et al. Massively parallel digital transcriptional profiling of single cells. Nat Commun. 2017;8:14049.

51. Butler A, et al. Integrating single-cell transcriptomic data across different conditions, technologies, and species. Nat Biotechnol. 2018;36(5):411–20.

52. Hafemeister C, and Satija R. Normalization and variance stabilization of single-cell RNA-seq data using regularized negative binomial regression. Genome Biol. 2019;20(1):296.

53. Street K, et al. Slingshot: cell lineage and pseudotime inference for single-cell transcriptomics. BMC Genomics. 2018;19(1):477.

54. Trapnell C, et al. TopHat: discovering splice junctions with RNA-Seq. Bioinformatics. 2009;25(9):1105–11.

55. Langmead B, and Salzberg SL. Fast gapped-read alignment with Bowtie 2. Nat Methods. 2012;9(4):357–9.

56. Lawrence M, et al. Software for computing and annotating genomic ranges. PLoS Comput Biol. 2013;9(8):e1003118.

57. Robinson MD, et al. edgeR: a Bioconductor package for differential expression analysis of digital gene expression data. Bioinformatics. 2010;26(1):139–40.

58. Granja JM, et al. ArchR is a scalable software package for integrative single-cell chromatin accessibility analysis. Nat Genet. 2021;53(3):403–11.

59. Prescott SL, et al. Enhancer divergence and cis-regulatory evolution in the human and chimp neural crest. Cell. 2015;163(1):68–83.

