## Supplemental Figures for "Integrated single cell analysis shows chronic alcohol drinking disrupts monocyte differentiation in the bone marrow niche"

### Supplemental Figure 1

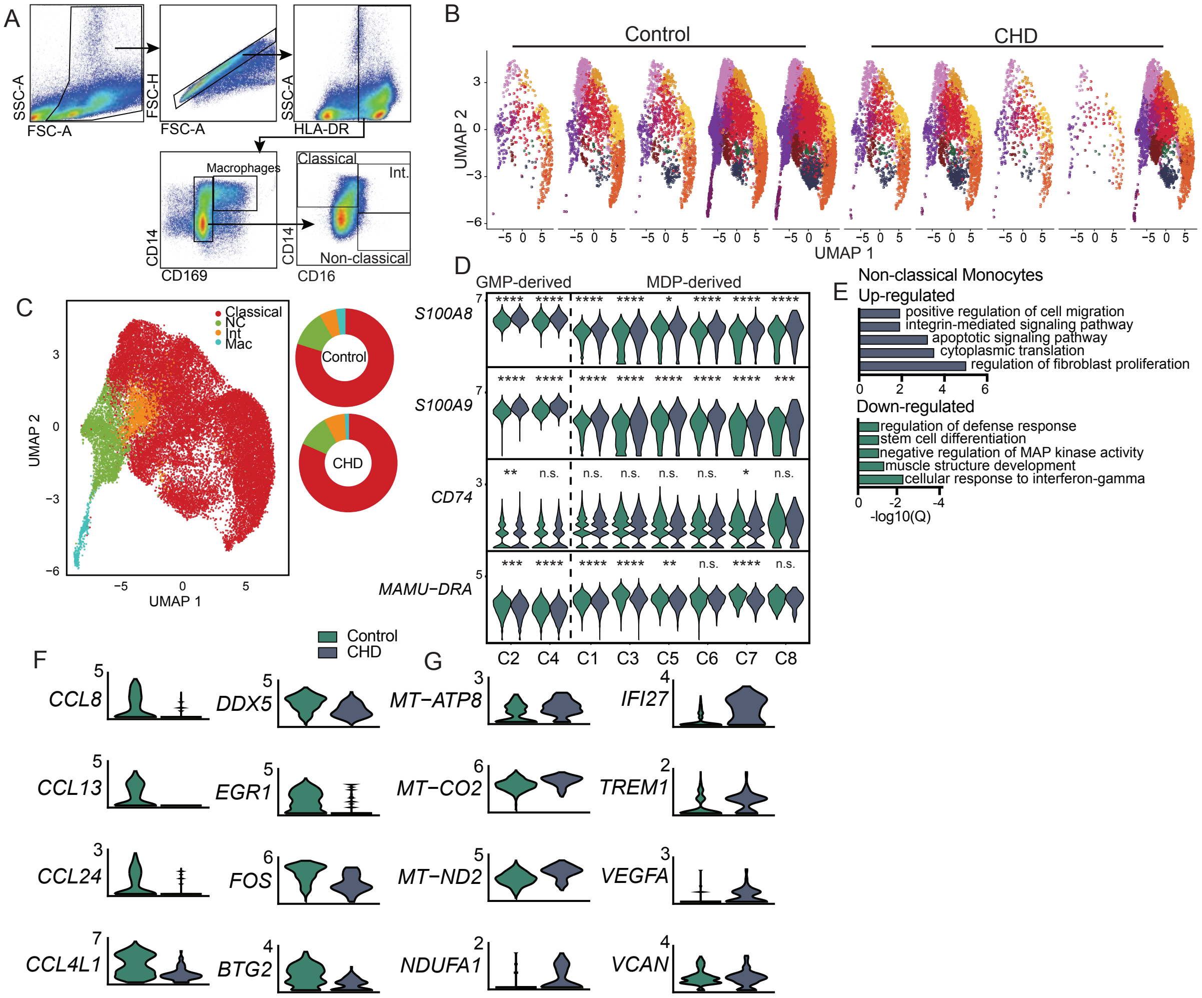


**Supp. Figure 1: scRNA-Seq of CD14+ cells from the bone marrow**

A) Example gating strategy for monocyte populations from the bone marrow. B) UMAPs of each individual macaque sample. C) UMAP annotated by broad classification of cell type and corresponding pie charts of abundance of cells from each group. D) Stacked violin plots of genes related to GMP or MDP lineages across each classical cluster and split by CHD and control groups. Statistical analysis performed by Mann-Whitney test where *=p<0.05, **=p<0.01, ***=p<0.001, ****=p<0.0001. E) Bar plot representing -log$_{10}$(q-value) functional enrichment scores for genes up- (blue, top) and downregulated (green, bottom) in non-classical monocytes with CHD. F,G) Violin plots of down- (F) and up- (G) regulated DEG with CHD in the macrophage cluster.

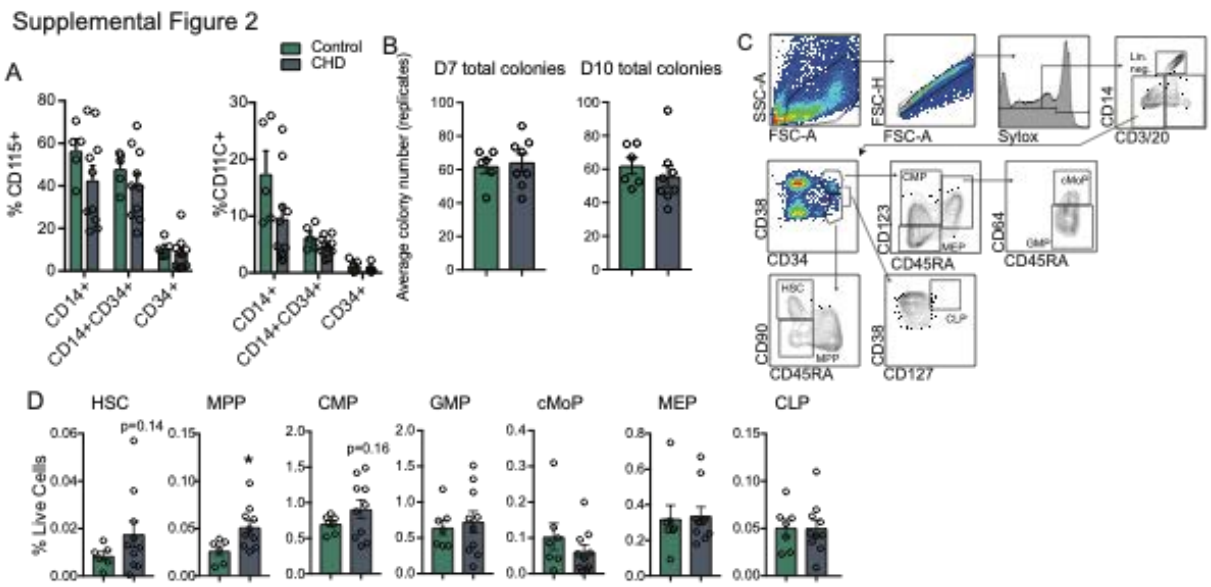


**Supp. Figure 2: CD34+ progenitor differentiation assays and flow cytometry of CD34+ cells from the bone marrow**

A) Bar plots showing percentage of CD115+ (left) and CD11C+ (right) in the indicated culture populations from flow cytometry. B) Bar graphs showing average colony numbers on day 7 and day 10 of culture. C) Gating strategy for bone marrow progenitor cells. D) Bar plots showing percentages of progenitor populations in the bone marrow compartment determined by flow cytometry. Unless indicated, statistical significance was tested by t-test with Welch's correction. *=p<0.05, **=p<0.01, ****=p<0.0001.

Supplementary Figure 3

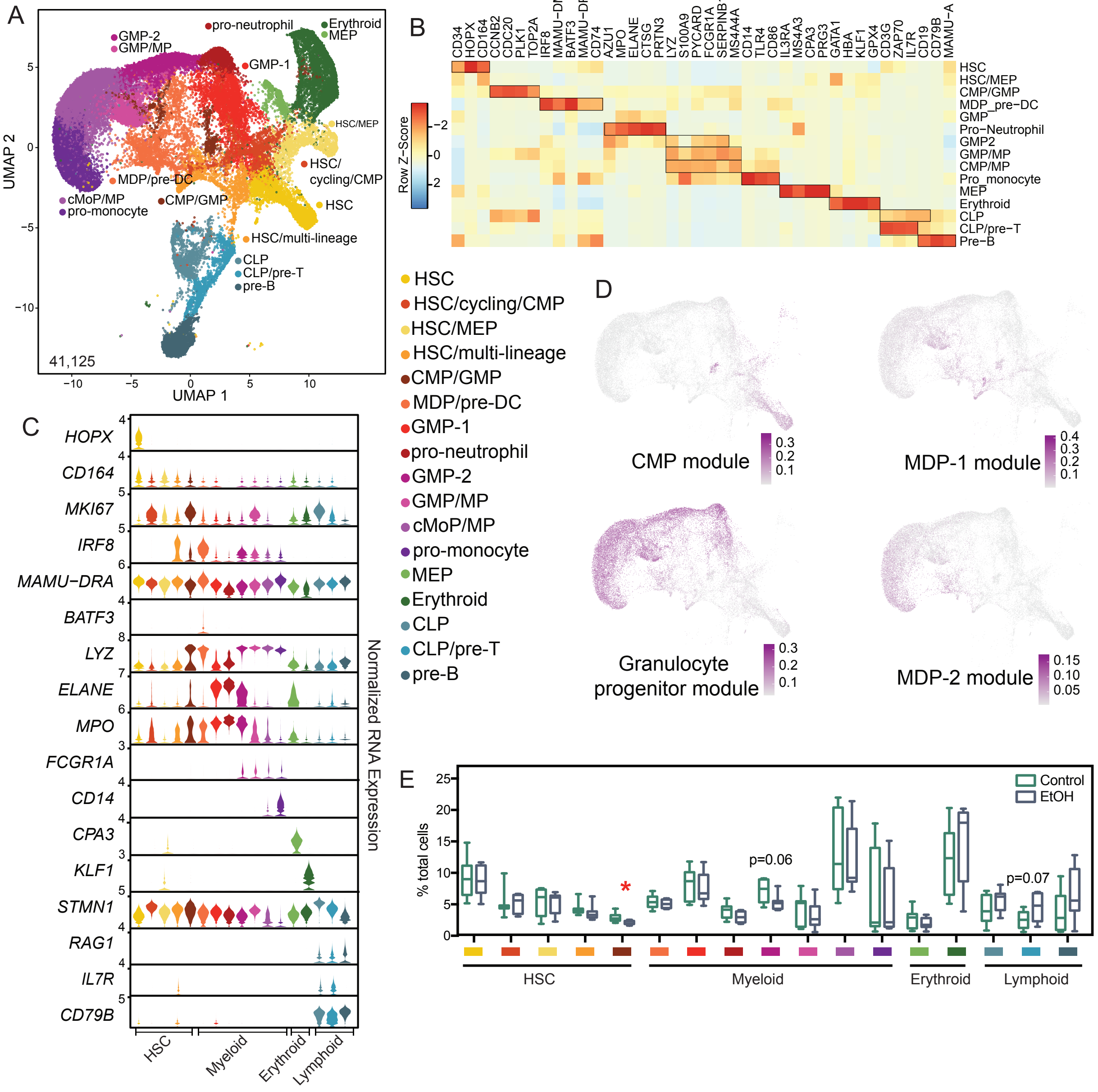

**Supp. Figure 3: scRNA-Seq of CD34+ cells from the bone marrow**

A) UMAP clustering of 41,125 cells and indicated cluster identification. B) Heatmap showing averaged gene expression of highly expressed genes from each progenitor cluster. C) Stacked violin plot showing expression of genes identified using Seurat's *FindAllMarkers* function. D) Feature plots showing relative expression of module scores for CMP, Granulocyte progenitors, MDP-1, and MDP-1 in the myeloid lineage. E) Box plots of percentages of each cluster across control and CHD groups. Statistical analysis performed by two-way ANOVA with multiple comparisons, *=p<0.05.

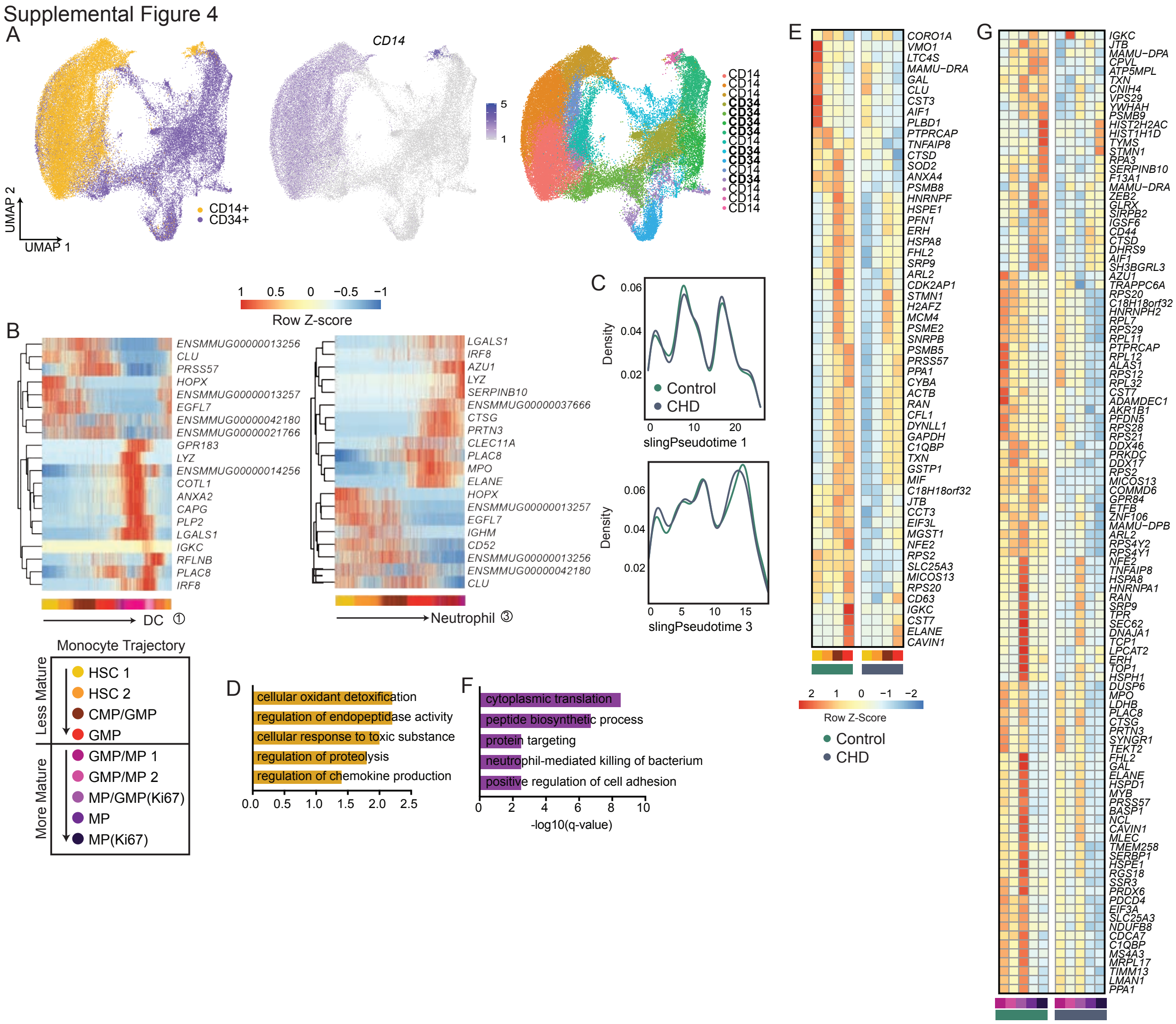

**Supp. Figure 4: Myeloid progenitor subsets in the bone marrow**

A) UMAP projection of integrated CD34+ (Figure 4) and CD14+ (Figure 2) datasets (left). Feature plot of *CD14* expression to identify contaminating CD14+ cells (middle). UMAP of Seurat clusters labeled with CD14 or CD34 designation for selection of only CD34+ cells (right). B) Heatmaps of genes explaining the DC (left) and neutrophil (right) lineage trajectories. C) Cell density plots for Control and CHD groups across the DC (top) and neutrophil (bottom) trajectory lineages determined by Slingshot. D,F) Bar plot showing functional enrichment terms from less (D) and more (F) mature clusters. E,G) Heatmap showing averaged gene expression of upregulated DEG from less (E) and more (G) mature clusters split by CHD and control groups.

Supplemental Figure 5

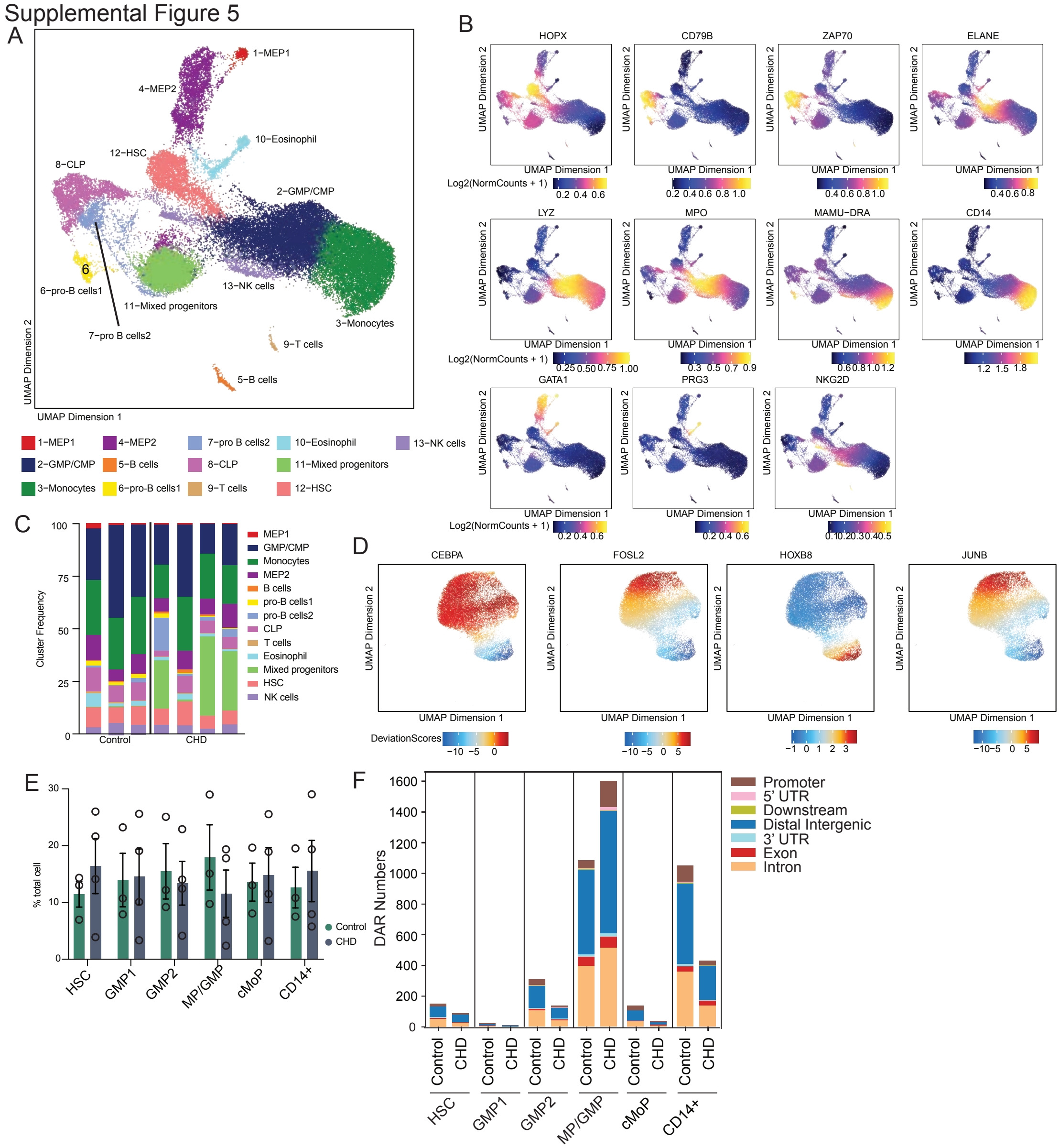


**Supp. Figure 5: scATAC-Seq of bone marrow CD34+ and CD14+ cells**

CD34+ and CD14+ cells were sorted from bone marrow and subjected to 10X scATAC-Seq. A) UMAP projection of all cells based on gene accessibility. B) Feature plots of indicated marker genes from the gene score matrix. C) Stacked bar plot distribution of clusters across each sample. D) UMAP plots of motif accessibility for each of the indicated transcription factors. E) Bar plot of percentages of control and CHD groups across each cluster. F) Bar plots of the percent of differentially accessible regions (left) and total numbers of differentially accessible regions (right) between control and CHD samples.

**Supp. Table 1: Animal cohort characteristics**

**Supp. Table 2: FindAllMarkers (CD14+, Culture, CD34+)**

**Supp. Table 3: Module Score Gene Lists**

**Supp. Table 4: Marker GeneScores scATACseq**
